# Super-resolved live-cell imaging using Random Illumination Microscopy

**DOI:** 10.1101/2020.01.27.905083

**Authors:** Thomas Mangeat, Simon Labouesse, Marc Allain, Emmanuel Martin, Renaud Poincloux, Anaïs Bouissou, Sylvain Cantaloube, Elise Courtaux, Elodie Vega, Tong Li, Aude Guénolé, Christian Rouvière, Sophie Allard, Nathalie Campo, Magali Suzanne, Xiaobo Wang, Grégoire Michaux, Mathieu Pinot, Roland Le Borgne, Sylvie Tournier, Jérôme Idier, Anne Sentenac

## Abstract

Super-resolution fluorescence microscopy has been instrumental to progress in biology. Yet, the photo-induced toxicity, the loss of resolution into scattering samples or the complexity of the experimental setups curtail its general use for functional cell imaging. Here, we describe a new technology for tissue imaging reaching a 114nm/8Hz resolution at 30 µm depth. Random Illumination Microscopy (RIM) consists in shining the sample with uncontrolled speckles and extracting a high-fidelity super-resolved image from the variance of the data using a reconstruction scheme accounting for the spatial correlation of the illuminations. Super-resolution unaffected by optical aberrations, undetectable phototoxicity, fast image acquisition rate and ease of use, altogether, make RIM ideally suited for functional live cell imaging *in situ*. RIM ability to image molecular and cellular processes in three dimensions and at high resolution is demonstrated in a wide range of biological situations such as the motion of Myosin II minifilaments in *Drosophila*.

## Introduction

Cell biology began with the light microscope in the seventeenth century. Since then, optical microscopy has remained an essential tool for cell biologists: learning how cells function requires a detailed knowledge of their structural organization and of the dynamic interplay of their many constituents, often over extended periods of time. A decisive breakthrough in microscopy was the specific tagging of virtually any protein with a fluorescent probe to visualize its location, dynamics, and potential interactions with other partners in living cells. But imaging subcellular structures required improving the resolution beyond the diffraction barrier, about 300 nm, which limits widefield microscopes. Furthermore, reaching the highest resolution attainable while using the least possible light intensity to preserve live cell integrity raised two challenges that seemed to be mutually exclusive.

In the past two decades, the development of super-resolution fluorescence imaging techniques have been developed to break the diffraction limit, notably stimulated emission depletion (STED) (Hell and Wichmann, 1994; Klar and Hell, 1999), stochastic optical reconstruction microscopy and photoactivated localization microscopy (STORM/PALM) (Betzig et al., 2006; Hess et al., 2006; Rust et al., 2006), or structured illumination microscopy (SIM) (Heintzmann and Cremer, 1999; Gustafsson, 2000; Gustafsson et al., 2008). These techniques and their later improved versions have provided impressive details of subcellular structures (for a review see Sahl et al., 2017). But each of them present caveats that limit their general use for live-cell imaging. Saturated fluorescence (STED), pointillists methods (STORM and PALM) and intrinsic fluorescence fluctuation approaches (Dertinger et al., 2009) reach their performance at the cost of intense light shining and/or prolonged data acquisition time that restrict imaging to small volumes of observation or slow temporal dynamics. Therefore, the vast majority of what is being looked at by super-resolution microscopy (SRM) is fixed cells with the possibility of sample distortions and artifacts induced by chemical treatments (Richter et al., 2017).

Currently, Structured illumination microscopy (SIM) presents the best compromise between spatial and temporal resolutions with low toxicity for live imaging. The SIM super-resolved image is formed numerically from several low-resolution images obtained for different positions and orientations of a periodic illumination pattern. The success of the numerical reconstruction relies on a precise knowledge of the illumination. When aberrations, possibly induced by the sample itself, distort the illumination pattern, the reconstruction fails. Thus, the best SIM resolution, about 120 nm transversally and 360 nm axially, is obtained with thin, transparent cell monolayers (Shao et al., 2011). Others versions of SIM have been introduced for imaging subcellular processes in thicker samples, but at a lower xyz resolution, 220 x 220 x 370 nm for the lattice light sheet version (Chen et al., 2014; O’Shaughnessy et al., 2019), and about 160 x 160 x 400 nm for the spot-scanning illumination (AiryScan) (Sivaguru et al., 2018).

Recently, it has been demonstrated theoretically (Idier et al., 2018) and experimentally (Mudry et al., 2012; Labouesse et al., 2017) that periodical or focused illumination in SIM could be replaced by totally uncontrolled speckles. Counter-intuitively, the low-resolution images obtained with unknown speckle illuminations could be processed into a sample image of better resolution than classical widefield microscopy. Potentially, speckle illumination appeared to be ideally suited for live cell imaging: ease of use (no lengthy monitoring of experimental drifts, no time-consuming calibration protocols when changing the sample, objective or wavelength), widefield configuration, low levels of energy transfer to the samples and an extremely simple experimental setup. Yet, the resolution was too low for imaging subcellular dynamics.

In the present work, we developed a technique based on speckle illumination, that we call Random Illumination Microscopy (RIM), which achieves a super-resolution level comparable to the best 3D periodic SIM, with the ease of use and application range of widefield microscopy. The gain of resolution was obtained using an original data processing combining the statistical approach of fluctuation microscopy with the demodulation principle of structured illumination microscopy. Most importantly, speckle illumination is insensitive to optical aberrations and scattering by thick specimens, and shows minimal phototoxicity, making RIM a method of choice for live-cell imaging *in situ*.

RIM ability to visualize biological processes in 3D, at high resolution and over extended periods of time is demonstrated, in comparison with the best available imaging techniques, on a wide range of macromolecular complexes and subcellular structures in action, such as the Z-ring of dividing bacterial cells, the dynamic actin network of macrophage podosomes, the mobility of proliferating cell nuclear antigen (PCNA) during DNA replication, or kinetochore dynamics in mitotic *S. pombe* cells. Analysis of multicellular samples was illustrated by imaging the intestine microvilli of *C. elegans*, the 3D motion of myosin minifilaments within developing *Drosophila* tissues, or the collective invasive migration of border cells in fly ovary. These examples illustrate the wide range of possible applications of RIM for imaging live cell functions and related pathologies *in vivo*. The simplicity of RIM experimental setup and operation mode should hopefully democratize super-resolution microscopy, at low cost, in cell biology laboratories.

## Results

### Principle of RIM

In RIM, a super-resolved reconstruction of the sample is formed numerically from several low-resolution images of the sample recorded under different uncontrolled speckle illuminations, hereafter named speckle images. A speckle is the light pattern formed by a coherent (laser) beam after the reflection or transmission by a random medium (Figure 1A). To implement RIM, a standard widefield epi-fluorescence microscope was modified by replacing the lamp with different laser diodes and introducing a Spatial Light Modulator (SLM), displaying random phase masks along the illumination path. Several hundreds of different speckles could be generated per second by changing the SLM display (Figure 1A-B; Movie S1).

**Figure 1.**
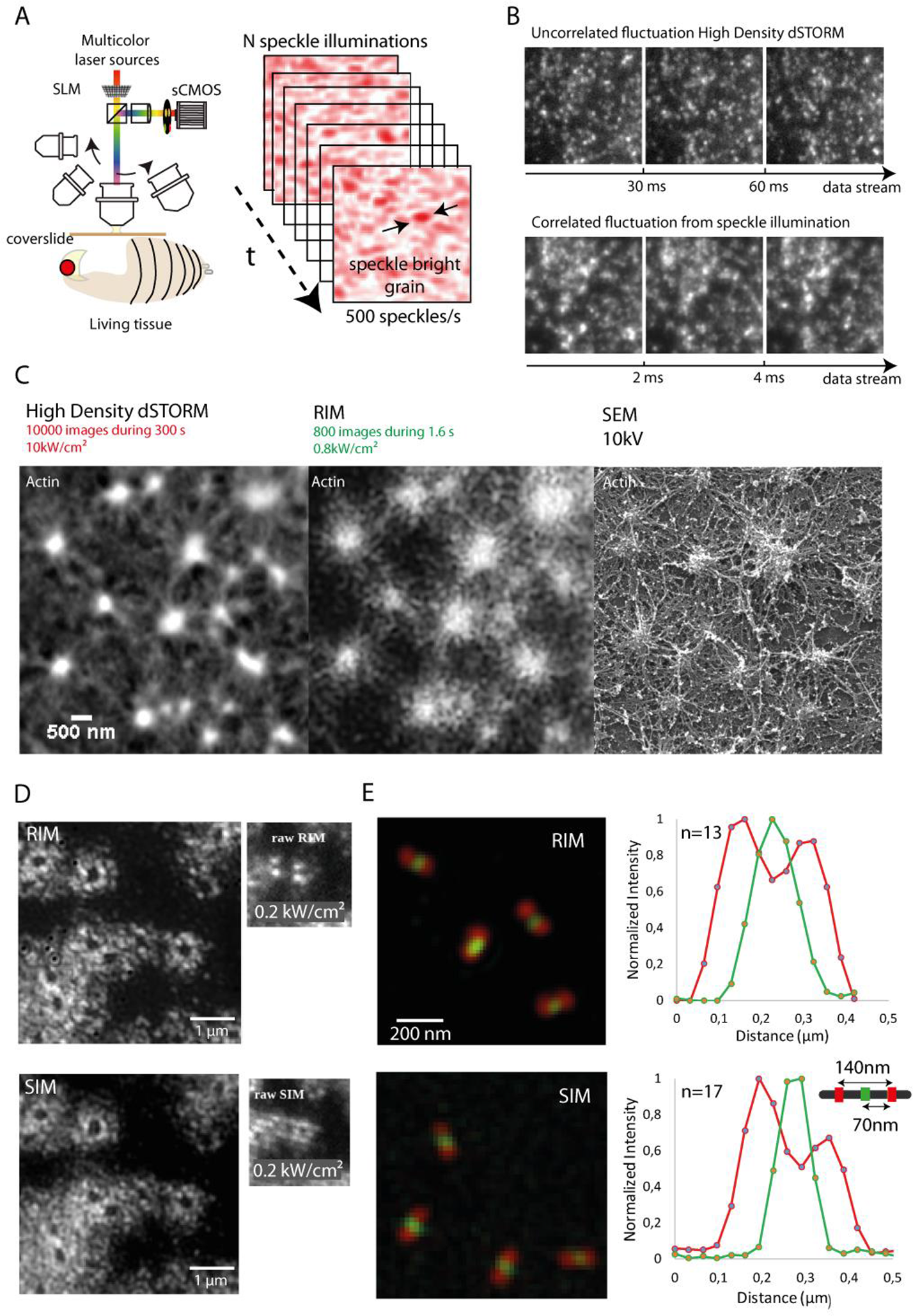
RIM *versus* fluctuation microscopy and Structured Illumination Microscopy. (A) RIM schematic implementation. A low cost Spatial Light Modulator (SLM) acting as a diffuser is implemented in a classical inverted microscope. Illuminated by multicolor lasers, the SLM sends five hundred different speckle patterns per second on the specimen. The fluorescence light is collected onto an sCMOS camera after appropriate filtering. The same set-up is operational for different objectives with numerical apertures between 0.15 and 1.49, and for wavelengths in the range of 450 nm to 600 nm. (B) Stream of raw data obtained with high-density fluctuation microscopy (dSTORM) or RIM (speckle illumination). One dSTORM image required 30 ms integration time while one RIM image required 2 ms. The sample is fixed macrophages, stained for F-actin, that have been unroofed to leave only the podosomes at the surface of the coverslip, shown in (C). (C) Super-resolved images of the macrophages F-actin network, obtained by processing 10,000 dSTORM raw images as in (B) using (Nanoj software SRRF; Gustafsson et al., 2016) or 800 raw RIM images using algoRIM, compared to a reference Scanning Electron Microscopy image. Note that RIM reconstruction is closer to the SEM image and does not suffer from the non-linear response to brightness of fluctuation microscopy. (D) Comparison between RIM and SIM imaging using the same experimental set-up. The RIM raw data are obtained by displaying 200 different random patterns on the SLM while the SIM data are obtained by displaying 30 different orientations and translations of a periodic pattern. Both experiments are performed with the same total number of photons injected into the sample (4 µJ per pixel). The sample corresponds to unroofed fixed macrophages, as in Figure 1C, with tagged vinculin surrounding the podosomes cores attached to the substrate. Right, comparison between RIM and SIM raw images. Left, RIM and SIM reconstructions. The dynamics and the resolution of the reconstructions are similar. (E) Comparison between RIM and SIM resolution using the commercial ELYRA microscope for two-color imaging. The sample is a calibrated DNA nanoruler (SIM 140 YBY), where two red fluorophores (Alexa 561) attached at the DNA ends are separated by 140 nm, and are equidistant (70 nm) to a green fluorophore (Alexa 488). Both RIM and SIM estimated the red-to-green distance to about 70 nm as evidenced in the graphs displaying the green and red intensities with respect to the distance averaged over n=13 and n=17 nanorulers, respectively, using a co-location analysis. The total RIM imaging process took less than 10 min from the insertion of the sample in the microscope to the reconstruction. In contrast, the total SIM imaging process took about 2 h for completing the calibration, acquisition and checking steps (Ball et al, 2015 and Demmerle et al, 2017). In addition, while three parameters needed to be tuned for the RIM reconstruction scheme, seven were required for the SIM inversion method.

The random bright grains of the speckle illumination, depicted in Figure 1A, ensure a quasi-pointillist excitation of the fluorescence, which connects RIM to the field of high-density fluctuation microscopy (dSTORM) (Baddeley et al., 2009), as illustrated in Figure 1B. Yet, in fluctuation microscopy, the excited fluorophores are sparser and appear at uncorrelated positions while in RIM they are excited collectively within the speckle bright grains over a typical distance of half a wavelength corresponding to the speckle correlation length. Due to this collective excitation, RIM requires significantly less images to cover the whole sample and a smaller integration time than fluctuation microscopy.

The super-resolved reconstruction is formed from the speckle images using a reconstruction scheme named AlgoRIM. AlgoRIM is based on a rigorous mathematical analysis (Idier et al., 2018) that takes advantage of the spatial correlation of the fluorophore excitation induced by the speckle to gain a two-fold increase in resolution. The fluorescence high frequency features are extracted from the variance of the raw images, as in fluctuation microscopy, through a demodulation process using the speckle autocorrelation as a carrier wave, as in Structured Illumination Microscopy. AlgoRIM does not invoke the sparsity of the excitation or the binarity of the fluorescence for achieving super-resolution and, even though it is based on the variance of the speckle images, it yields a linear response to brightness (for more details, see Supplemental information).

In Figure 1C, we compare the reconstructions of the same sample of tagged F-actin network in podosomes obtained by second-order statistics dSTORM (*NanoJ software, Super Resolution Radial Fluctuation*) (Gustafsson et al., 2016) and RIM. Podosomes are actin-rich, cell adhesion structures applying protrusive forces on the extra cellular environment, recently observed by 3D dSTORM (Bouissou et al., 2017). Remarkably, the RIM reconstruction showed more details than dSTORM at the level of podosomes nodes, and was closer to a Scanning Electron Microscopy (SEM) image of a similar sample. Importantly, RIM required ten times less images, used ten times less power and was two hundred time faster than dSTORM.

In Figure 1D, we compare the reconstructions and the raw images of the same sample of vinculin-tagged podosomes attached to the substrate obtained by RIM and SIM (by displaying either random or periodic masks on the SLM). The resolution and overall dynamic range of the two techniques are remarkably similar. To evaluate more precisely the resolving power of RIM, we imaged a calibrated DNA nanoruler (SIM 140 YBY), where two red fluorophores (Alexa 561) attached at the DNA ends are separated by 140 nm, and are equidistant (70 nm) to a green fluorophore (Alexa 488). In Figure 1E, we compare the RIM reconstruction to the SIM image given by the commercial two-color Zeiss SIM Elyra system. Figure 1E shows that both RIM and SIM succeeded in separating the red fluorophores and accurately located the green middle one. RIM resolution was the same of that of SIM with an average red-to-green distance of about 70 nm in both cases (compare the graphs in Figure 1E). Additional experiments with calibrated samples indicated that RIM resolution matched that of the best periodic SIM techniques, about 120 nm for fluorophores emitting at 514 nm with an objective of NA=1.49 (Figures S1E and S3A).

In these types of experiment where the sample does not distort the illumination pattern, the major interest of RIM compared to SIM is the extreme simplicity of the experimental protocol which can be performed in less than ten minutes even for two-color imaging. Since the illumination patterns do not need to be known, the only tuning required before imaging consists in checking the focus. In contrast, in the case of SIM, the knowledge of the illumination patterns is mandatory, which implies a specific sample preparation and a precise microscope alignment and polarization control for the two colors, which, altogether, may take about two hours (Demmerle et al., 2017). Another interest of RIM, compared to SIM, is the robustness and ease-of-use of its inversion procedure. AlgoRIM required the tuning of 4 parameters, the widths of the observation point spread function and speckle correlation and two Tikhonov parameters (see Supplemental information), while at least 7 were needed for the SIM reconstruction procedure, in particular for recovering the illumination patterns from the raw images. This last task is particularly delicate as it can be jeopardized by a too big difference between the excitation and fluorescence wavelengths (Figure S1F).

### RIM allows high fidelity super-resolved imaging in three-dimensions

RIM three-dimensional (3D) imaging is obtained by translating the sample through the focal plane (Movie S1) and by recording several speckle images at each position. The speckle illumination and the variance-based data processing ensure an efficient optical sectioning (Ventalon and Mertz, 2005; Ventalon et al, 2007). Note that, when necessary, the spherical aberration induced by the index mismatch between the objective immersion oil and the mounting medium has been accounted for when reconstructing the 3D image (Sibarita, 2005).

To test the fidelity of the super-resolved images obtained with 3D RIM, we focused on dense filamentous structures. We imaged the vimentin network from fixed HUVEC cells and reconstructed the whole network from 200 speckles per slice, 12 slices and an axial step of 100 nm (see in Figure 2A the color-coded axial position of the filaments, and Movie S2). As seen in Figure 2B, RIM transverse resolution was much better than that of confocal microscopy and similar to that of STED microscopy, about 120 nm (with fluorophores emitting at 700 nm and a NA of 1.49). Interestingly, RIM reconstruction was free from common artefacts such as the disappearance or merging of filaments (Marsh et al, 2018) and it provided the same image as STED both in the dense and sparse regions of the sample. A total of 1 kW/cm², five times less than that required for confocal microscopy, was delivered to the entire volume (30 µm x 30 µm field of view) in less than 3 seconds.

**Figure 2:**
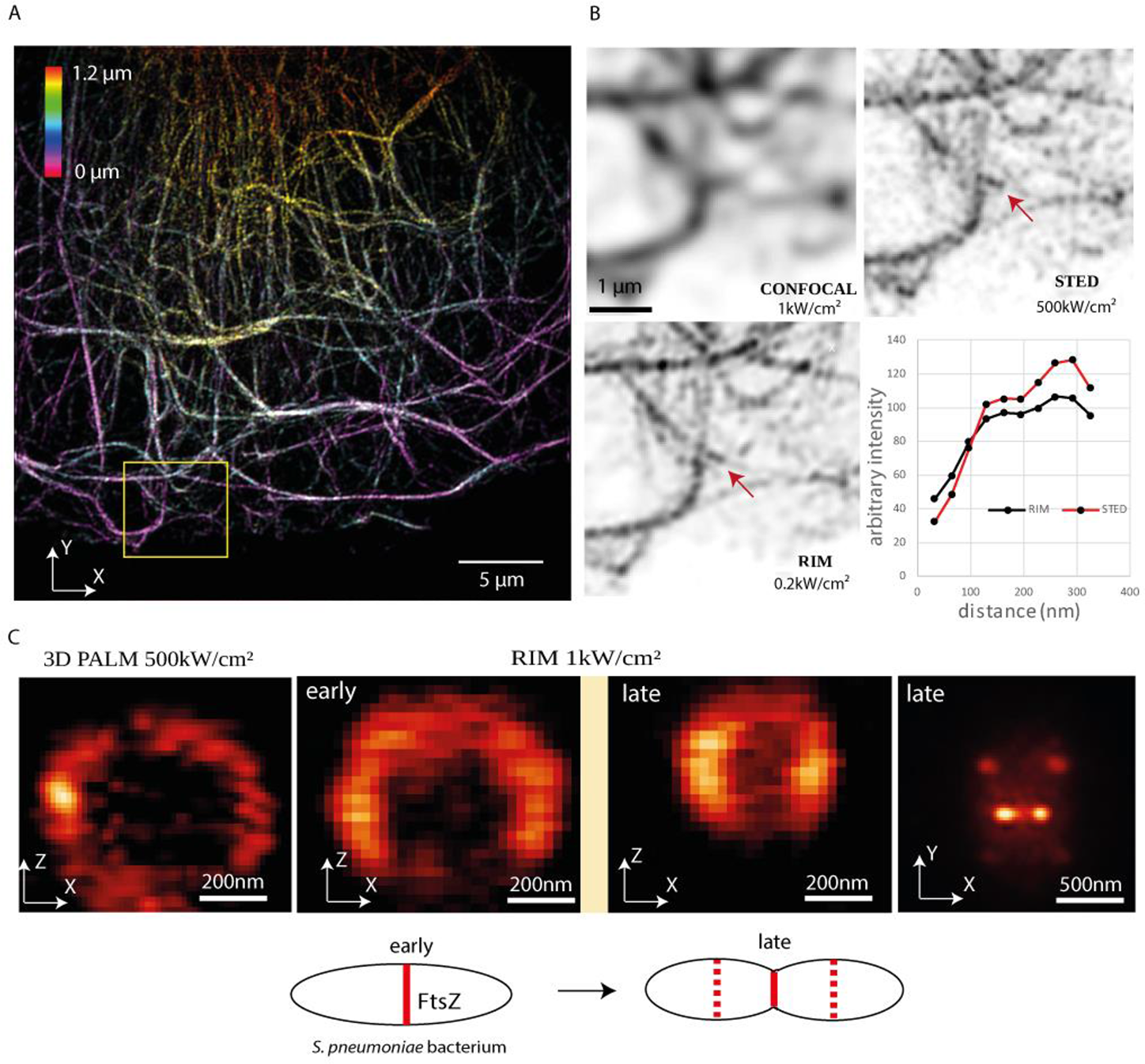
**RIM allows high-fidelity super-resolved live imaging in the three dimensions.** (A) RIM 3D view of dense vimentin network from fixed HUVEC cell using a fluorescent antibody dedicated for STED microscopy with emission at 700 nm. The 3D image is made of 12 slices 100 nm apart. The color scale indicates the axial position (Fiji image processing). The yellow square locates the filaments enlarged in (B). (B) The same vimentin filaments were observed by confocal microscopy (1 kW/cm²), STED microscopy (500 kW/cm²) or RIM at low photon budget (200 W/cm²). The curves depict the intensity recorded along the two dots shown by the arrow in RIM and STED images. (C) Axial and transverse cuts of 3D RIM images of *S. pneumoniae* Z-rings containing FtsZ tagged with mEos3.2 taken at different stages of the cell division compared to 3D PALM equipped with adaptive optics. The 3D image is made of 20 slices, 64 nm apart. Early and late FtsZ annular constrictions from two dividing cells are shown (Z-ring diameters of 640 nm or 400 nm) in the axial cut. Right, transverse cut at the equatorial plane of the Z-rings of the two attached daughter cells (late division stage).

To further investigate the axial resolution, we tested the ability of RIM to resolve the cell division ring of the bacteria *Streptococcus pneumoniae.* Cells were labeled with fluorescent FtsZ, an homolog of tubulin that can polymerize and assemble into a ring, called the Z-ring, at the site were the septum forms during cell division. This ring whose diameter ranges from 0.3 to 0.9 µm depending on the division stage (Fleurie et al., 2014) is roughly perpendicular to the observation focal plane when the bacteria lies on the substrate, which makes it ideally suited for checking RIM axial resolution. Z-rings were imaged in live bacterial cells transferred in minimum medium using 3D-PALM or RIM (Figure 2C). When comparing RIM and PALM images, note that the FtsZ protein was tagged differently to allow for the specific requirements of PALM. This may affect differently the level of expression of FtsZ and the efficiency of Z-ring formation. RIM image reconstructions clearly revealed the heterogeneous structure of Z-rings formed of discontinuous clusters of FtsZ protein, with highly different levels of fluorescence intensity, though at a lesser resolution than PALM equipped with adaptive optics (Zheng et al., 2017). Interestingly, in the transverse cut at the equatorial plane of a late dividing cell, significant levels of FtsZ protein were observed in-between the two nascent Z-rings on each side of the central ring that disassembles as it constricts, suggesting a constant exchange of free and ring-associated FtsZ molecules. The resolution of RIM was estimated to be 120 nm transversally and 300 nm axially, which is equivalent to that of the best SIM (Fleurie et al., 2014).

### RIM provides super-resolved movies of live specimens at high temporal resolution and low toxicity

To optimize the temporal resolution of RIM and limit phototoxicity, one should know the minimal number of raw speckle images that is necessary for a faithful reconstruction of a given specimen. The latter depends on the nature of sample (dense or sparse) and its dynamics. Tests on fixed samples showed that super-resolved images can be obtained with only 50 speckles but at the cost of a residual illumination inhomogeneity (Figure S3A). Increasing the number of speckles improves the illumination homogeneity and the signal to noise ratio but movements of the live specimen during the recording may blur the reconstruction (Figure S3B). The speckle images being recorded regularly during the observation time, different trade-offs can be tested with the same data set. In addition to a classical processing in which the total number of speckle images recorded during the experiment, T, is divided into stacks of N speckle-images for forming T/N super-resolved reconstructions, we considered an interleaved reconstruction scheme in which the stacks of N speckle images are shifted by Q images (with Q<N) to form T/Q super-resolved reconstructions as illustrated in Figure 3B. The interleaved strategy permitted to improve the temporal resolution (by diminishing Q) while keeping a good signal to noise ratio and illumination homogeneity (using large enough N). The optimal choice of Q and N may vary depending on the sample.

**Figure 3:**
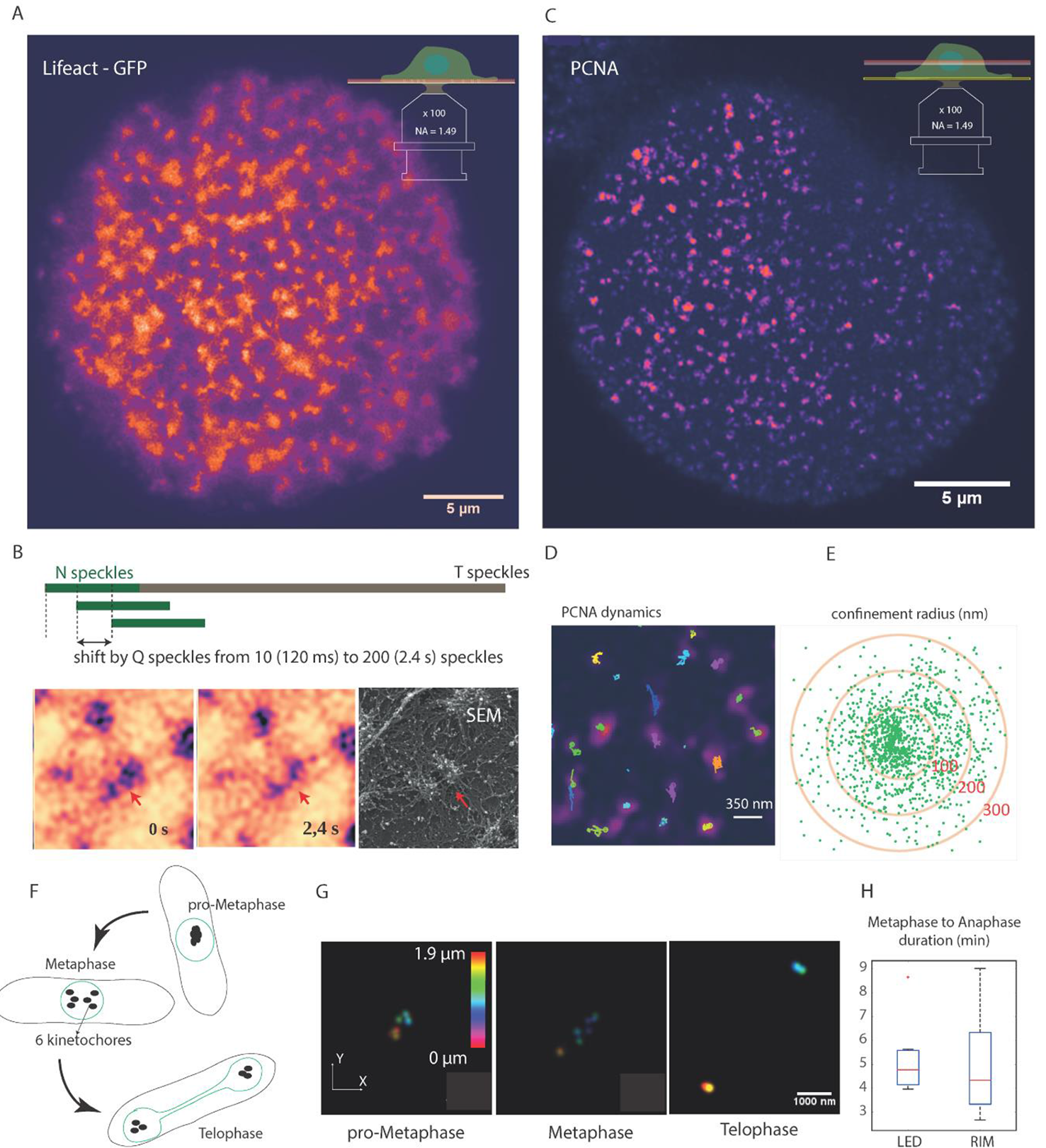
**RIM provides images with 120 nm resolution of dense and sparse samples with an adjustable temporal resolution and phototoxicity compatible with live imaging.** (A) 20 min dynamic imaging of human macrophage podosomes labeled with Lifeact-GFP, (frame from Movie S3). The temporal resolution was 0.12 s with an interleaved reconstruction strategy based on sliding windows of 800 speckles for image reconstruction. (B) Top: Principle of the interleaved reconstruction allowing an adjustable temporal resolution. A total of T (typically 100,000) speckle images are recorded every 12 ms for the whole movie. Stacks of N (typically 800) raw: images shifted every Q (typically 10) images are used for forming the super-resolved reconstruction. The N and Q numbers are adjusted depending on the nature of the sample (dense or sparse) and the time-scale of the biological events. Bottom: evolution of F-actin core podosome cores observed with RIM and compared to a Scanning Electron Microscopy image. The filament linking the two nodes is clearly visible on the two images (red arrows). (C) Frame from Movie S4: PCNA dynamics during S phase in U2OS cell at 2 µm depth from the cover slide with a temporal resolution of 12.5 ms. (D) Trajectories of individual spot of PCNA during 20 s. Two families of trajectories can be observed. (E) Confinement of PCNA clusters. The average confinement distances of the slow and fast replication clusters are equal to 120 nm and 300 nm, respectively. (F) Schematic representation of fission yeast kinetochore displacement during mitosis. (G) Kinetochores harboring GFP-tagged Ndc80 protein were resolved in pro-metaphase, metaphase, and telophase. The color coded bar indicates the axial position (*S. pombe* typically measures 3 to 4 µm in diameter and 8 to 16 µm in length). (H) Statistical comparison of metaphase to anaphase duration as observed by RIM or classical widefield microscopy with synchronized LED illumination on 7 mitosis. The red segments indicate the median value, the box edges are the 25 and 75 percentiles and the whiskers extend to the most extreme datapoints.

To illustrate the different reconstruction strategies, we imaged the dense actin network of podosomes from human macrophages under live conditions (Figure 3A and Movie S3). Podosomes are composed of an approximatively 500 nm high and large F-actin protrusive core surrounded by an adhesion ring. The observation of their continuous spatial reorganisation requires super-resolved imaging over tens of minutes with a sub-second temporal resolution.

The compromise for imaging podosomes was to use stacks of 200 speckle-images in the classical reconstruction scheme (Figure S3C, Movie S3). We also implemented the interleaved strategy where stacks of 800 speckle images shifted by 200 to 10 images were used to form movies with temporal resolutions from 2.4 s to 0.12 s (Figure S3D, Movie S3). The 120 nm resolution of the RIM movie revealed actin filaments linking two actin cores, in agreement with observations by electron microscopy (Figure 3B). The robust estimation of podosome surpassed that obtained with live-SIM (van den Dries et al., 2019; Meddens et al., 2016) or other computational methods (Marsh et al., 2018). Notably, the photobleaching and toxicity of RIM was comparable to that of SIM (Figure S4D) and podosome dynamics could be observed during 20 min without detectable alteration of their reorganization.

A second example was the dynamics of the Proliferating Cell Nuclear Antigen (PCNA) (Figure 3C and Movie S4). PCNA is a protein involved in DNA replication, DNA repair, chromatin remodeling, and cell cycle. Here, the temporal resolution needs were higher, but the sparsity of the sample allowed us to use only 150 speckles in an interleaved reconstruction strategy yielding a temporal resolution of 0.12 s. The nanoclusters of PCNA were similar to those observed in fixed samples (Zessin et al., 2016). The trajectories of individual spots of PCNA were recorded during 20 s during the S phase of U2OS cells (Figure 3C, 3D and Movie S4). PCNA clusters mobility exhibited a fast and a slow diffusion regime, suggesting the existence of at least two pools of PCNA molecules probably belonging to different functional macromolecular complexes.

The average confinement distance of the slow replication clusters was about 120 nm while the fast diffusing PCNA clusters had a confinement about 200-300 nm (Figure 3E). These global results are in agreement with those previously obtained by single-particle tracking (Zessin et al., 2016). This experiment demonstrates the versatility of RIM, which, from the same set of data, provides both super-resolved images of the whole cell nucleus at all the phases of DNA replication and a trajectory analysis of PCNA similar to that obtained by single-particle tracking.

To illustrate the three-dimensional imaging potential and low toxicity of RIM, the mitosis of the fission yeast *Schizosaccharomyces pombe w*as recorded in a 3D movie with a temporal resolution of 20 s (Movie S5). *S. pombe* is a rod-shaped, symmetrically dividing eukaryotic cell that splits by medial fission. *S. pombe* possesses three chromosomes which can be tracked by live imaging during mitotic progression. Mitotic chromosomes are captured by microtubules at kinetochores, which are giant protein complexes assembled at the centromere of each sister chromatids. Kinetochores, fluorescently labelled on the Ncp80 protein, begin to align at the spindle center in phase 1 (spindle size of about 0.5µm; Mary et al., 2015). Remarkably, RIM imaging was able to visualize in xyz the 6 kinetochores attached to the three sets of sister chromatids in pro-metaphase (Figure 3G). In phase 2, the spindle retained roughly the same length, and the kinetochores oscillated between the two spindle pole bodies. In telophase, it was possible to observe the 3 kinetochores moving at each cell pole (Figure 3G). Importantly, this level of kinetochore resolution in live fission yeast cells has never been attained in the past. We also noted that the metaphase to anaphase duration under RIM illumination (4 min) was as expected (Figure 3H, Movie S5) indicating that the cells were not detectably stressed by the repeated speckle illuminations.

### RIM super-resolved imaging of optically aberrant and scattering tissues

A major bottleneck of a super-resolution imaging technique is its ability to keep its resolution level in optically aberrant or scattering environments.

We used *C. elegans* as a model multicellular organism to test RIM performance for imaging complex tissues like the worm intestine. A very high numerical aperture (TIRF objective with NA= 1.49) was selected for reaching the best possible resolution. In this configuration, aberrations are important due to the optical index mismatch between the worm and the mounting medium and the aberrations of the objective itself. In spite of this, RIM could image fluorescent ERM-1/ezrin, a protein constitutive of microvilli (microvilli are membrane protrusions that increase the surface area) and could clearly reveal the periodic organization of the intestine of living L4 larvae, at 15 µm depth (Figure 4A). The periodicity of microvilli was about 120 nm (Figure 4C) as confirmed by TEM images (Figure 4B).

**Figure 4:**
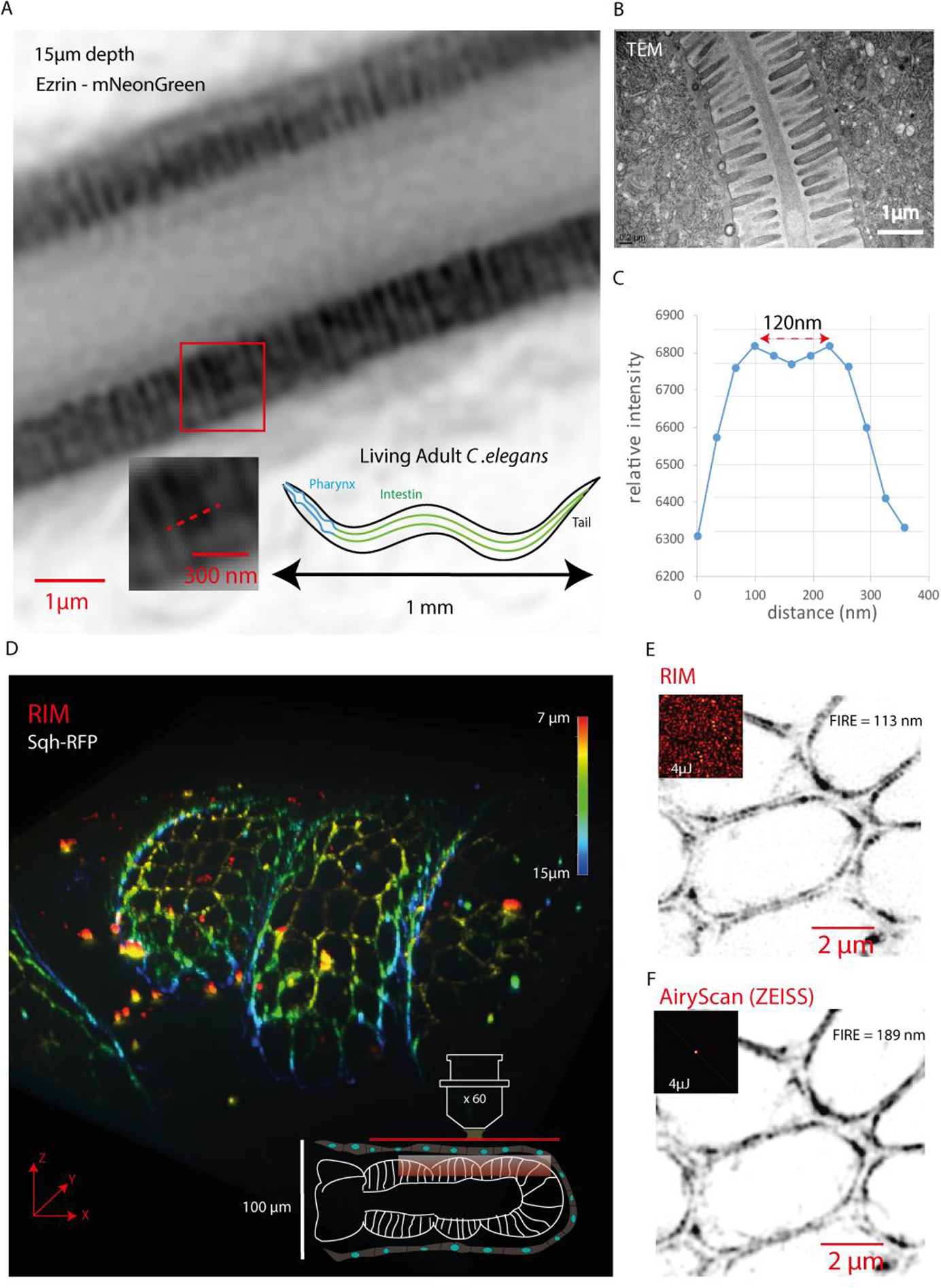
**RIM imaging of optically aberrant tissues without adaptative optics.** (A) RIM imaging of ERM-1/ezrin endogenously tagged with mNeonGreen discloses the microvilli brush border of the intestine of a live L4 larva (*C. elegans*). (B) Transmission electron microscopy (TEM) image of microvilli from L4 Larva (*C. elegans*). (C) The intensity profile between two microvilli, indicated by the red bar in (A), shows that they are separated by about 120 nm as confirmed by the electron microscope image (B). (D) RIM 3D imaging (the color codes for the axial position) of a large field of view (110µm x 110µm) of the RFP-tagged regulatory light chain of the non-muscle Myosin II motor protein, spaghetti squash (Sqh) at the apical plane of the epithelium of a fixed developing *Drosophila melanogaster* leg (inset). The acquisition of each plane took 400 ms (2 ms per speckle, 200 speckles). In comparison, the AiryScan technology needed 4.3 s for imaging the same field of view. (E-F) RIM (E) and Airyscan (F) images of the same sample. RIM and AiryScan Fourier Image Resolution (FIRE) resolution was estimated using Fourier Ring Correlation (FRC) technique (113 nm and 189 nm, respectively). The insets depict the illumination patterns in RIM (speckle) and AiryScan (focused spot). The same number of photons corresponding to 4 µJ per diffraction limited pixel was injected in the sample for RIM and AiryScan (see Figure S4).

In a second example, the tagged regulatory light chain of the non-muscle Myosin II motor protein, spaghetti squash (sqh), was imaged in a fixed developing leg of the fly *Drosophila melanogaster* (Figure 4D). Non-muscle Myosin II (thereafter referred to as Myosin II) is the major molecular motor generating contractile forces within non-muscle cells. The developing leg is a tissue 110 µm deep composed of a cylindrical columnar epithelium surrounded by a thin squamous epithelium (inset Figure 4D and Figure S4B). We tested RIM ability to produce super-resolution 3D images of Sqh-RFP in different regions of the apical plane of the columnar epithelial cells (Figure 4D). The squamous epithelium and the presence of numerous interstitial lipid droplets were important source of scattering and aberrations which made imaging difficult. Hence, this sample could not be imaged with periodic SIM because of the frequent disappearance of the illumination grid (see Figure S4C and Movie S6). The RIM images were thus compared to that obtained with the more robust focused scanning SIM known commercially as AiryScan. RIM images were better resolved than that of AiryScan (Figure 4E-F, Movie S7), and Fourier Ring Correlation (FRC) (Banterle et al., 2013) estimated a 113 nm Fourier Image Resolution (FIRE) resolution (Nieuwenhuizen et al., 2013) for RIM compared to 189 nm FIRE for AiryScan. This difference in resolution enabled RIM to distinguish the myosin dots aligned on the actin cortical networks of the cells. Imaging over a field of view of 110 µm x 110 µm required 0.4 s for RIM and 4.3 s for AiryScan. In addition, a careful investigation on several samples showed that, for the same energy (4 mJ) injected per voxel, RIM induced significantly less bleaching than AiryScan (Figure S4D).

### RIM multiscale imaging, from molecular motion to cell migration

To investigate the ability of RIM to visualize macromolecular motions deep inside several living tissues, we focused on Myosin II dynamics. Non-muscle Myosin II molecules are heterohexamers composed of two heavy chains, two regulatory light chains (Sqh), and two essential light chains. Myosin II hexamers assemble in an antiparallel manner to form 300 nm long minifilaments, comprising about 15 Myosin II dimers, that can be labeled at both ends with Sqh-RFP, forming a characteristic fluorescent doublet (Figure 5A), (Hu et al, 2017). Up to now, doublets of Myosin II minifilaments have only been observed in cultured cells (Beach et al, 2014; Fenix et al., 2016; Hu et al., 2017). In this work, we imaged Myosin II in different live *Drosophila* epithelia. Epithelial cells harbor three main pools of Myosin II, (Figure 5B II), two located apically called the junctional Myosin II accumulating at cell-cell adhesive contacts and the medial myosin inside the cell. A third pool is located basally at focal adhesion, and faces the basement membrane.

**Figure 5.**
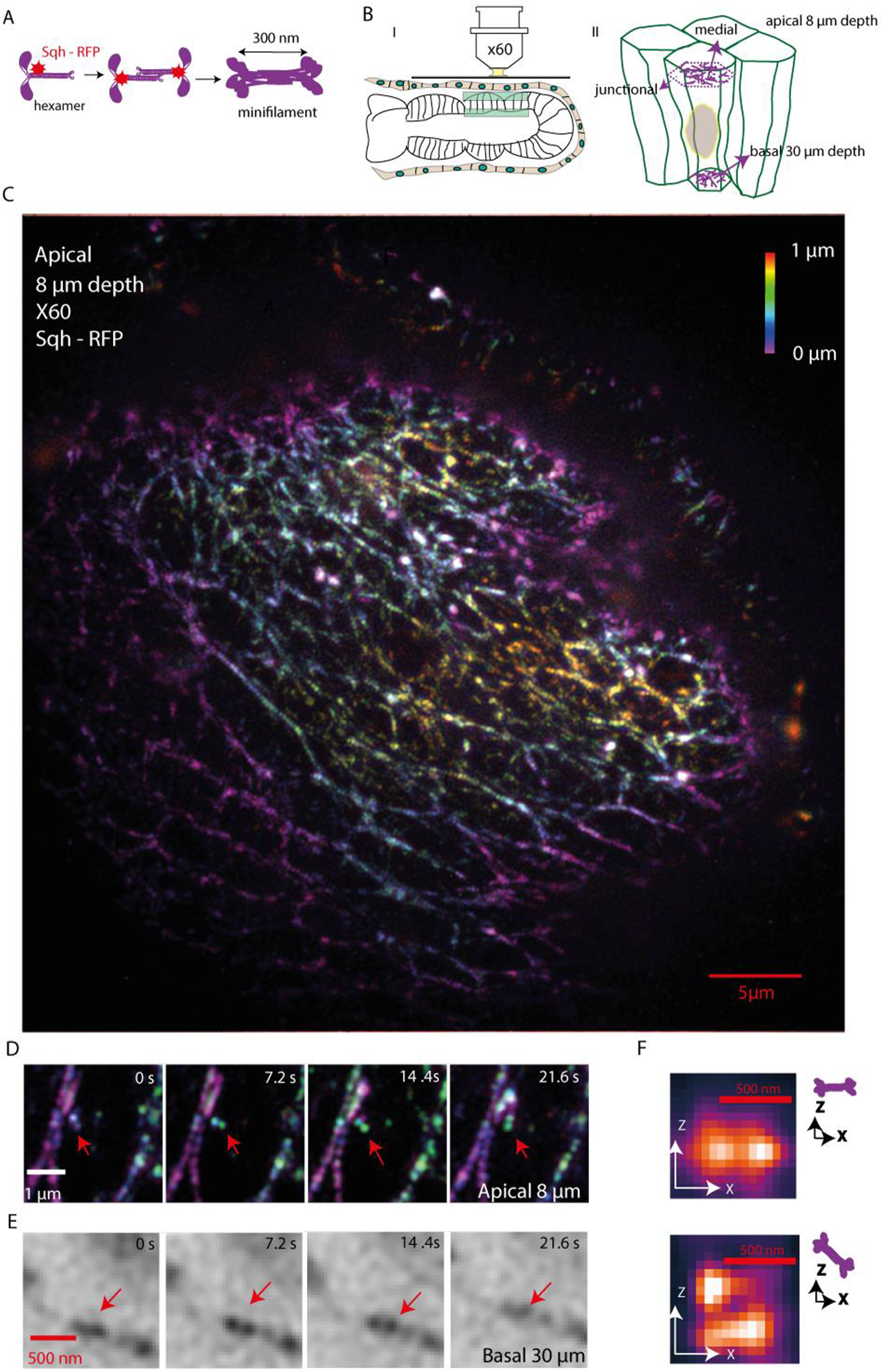
**3D RIM imaging of Myosin II in live developing Drosophila leg epithelium.** (A) Schematical representation of the assembly of non-muscle Myosin II minifilaments. The hexameric Myosin II protein, composed of two heavy chains, two essential light chains and two regulatory lights chains (Sqh-RFP) interact in an antiparallel manner to form dimers. About 15 dimers of Myosin II filaments are organized in parallel bundles lying along actin filaments. Myosin II molecules assemble into 300 nm-long bipolar minifilaments with Myosin heads symmetrically located on each side. *(B) (I)* Experimental conditions during acquisition; *(II)* Cartoon depicting the distribution of medial, junctional and basal pools of Myosin II filaments in *Drosophila* polarized epithelial cells. (C) RIM 3D widefield view of the regulatory light chain of Myosin II (Sqh-RFP) networks at the apical plane (8 µm depth) of *Drosophila* pupal leg. (D) Zoom illustrating the 3D motion of a single bipolar minifilament (red arrows) at the apical plane. (E) Zoom on the basal Myosin network taken at 30 µm depth illustrating the ability of RIM to follow a single bipolar minifilament deep in optically aberrant tissue (red arrows). (F) Super zoom showing the ability of RIM to distinguish the fluorescent doublet of a Myosin II minifilament whatever its orientation in the (x,z) plane (bicubic interpolation of the 3D image made of seven slices 150 nm apart).

We first imaged the basal plane of follicular epithelial cells (FEC) of *Drosophila* egg chamber at stage 9 where the imaging conditions, 1µm deep from the coverslip, are benign (Figure S5A-B). Myosin II filaments are organized in parallel bundles lying along actin filaments (Figure S5F) (He et al., 2010). Aligned fluorescent spots of Myosin II (Sqh-RFP) were well observed in the RIM widefield view (Figure S5C taken from Movie S8) and their motion, correlated by pair, confirmed their doublet nature (Figure S5D). In Figure S5E, two-color RIM imaged labeled Myosin II (Sqh-RFP) together with labeled Actin (Utrophin-ABD-GFP) and disclosed the interpenetration and alignment of Myosin and Actin filaments as expected (see the high magnifications in S5E, II and III).

For challenging the ability of RIM to visualize Myosin II molecular motion deep into a complex living tissue, we turned to the developing Drosophila leg whose imaging difficulty has already been pointed out in Figure 4D. Figure 5C shows a RIM 3D widefield view of the Myosin II (Sqh-RFP) network at the apical plane, 8 μm deep inside the live developing leg. RIM successfully visualized the pool of Myosin II concentrated at cell-cell contacts (junctional Myosin II). A zoom on this junctional Myosin revealed the distinctive spots corresponding to Myosin heads and showed the ability of RIM to follow the 3D motion of a single minifilament during 20 seconds (Figure 5D), independently of its orientation in the xyz directions (Figure 5F). Remarkably, single minifilaments could also be imaged at the basal plane of the epithelial cell, at 30 µm depth, despite the optically aberrant environment (Figure 5E). To our knowledge, no other imaging technique could follow, at such depths within a live tissue, the 3D displacement of myosin with such a resolution.

For a global functional analysis of Myosin II dynamics, taking advantage of the high spatio-temporal resolution of 3D RIM, we looked at the dorsal thorax of *Drosophila* pupal notum epithelium (Figure 6A). It is a single layer epithelium composed of epidermal cells and sensory organ precursor cells (SOP). Figure 6C shows a 3D RIM widefield view of the Myosin II networks at the apical plane, 7 μm from the coverslip. RIM clearly detected the two distinct pools of Myosin II, the medial Myosin II forming irregular networks inside the cells, and the junctional Myosin II accumulating neatly at cell-cell contacts. Each cell having its own pool of junctional Myosin II at the adhesive contacts, the 3D RIM image shows two parallel dotted lines corresponding to minifilaments myosin heads, not regularly spaced though in contrast to some previous observations on other fixed epithelial cells (Ebrahim, 2013). The fluorescence intensity plot between these two junctional myosin layers (red arrowheads in Figure 6C) indicates a 130 nm resolution. Even though this type of tissue was highly optically heterogeneous, as schematically presented in Figure 6A, the resolution remained constant (130 nm) over the whole field of view and was able to distinguish the Myosin filament fluorescent doublets (see Movie S8). Remarkably, RIM high temporal resolution was able to visualize the pulses of the medial Myosin II networks in a constant state of spatial reorganization (Martin et al., 2009) (Figure 6D, Movie S8). RIM was also able to visualize the flow of junctional myosin II at the cell cortex (Rauzi et al., 2010) (Figure 6E, Movie S8).

**Figure 6.**
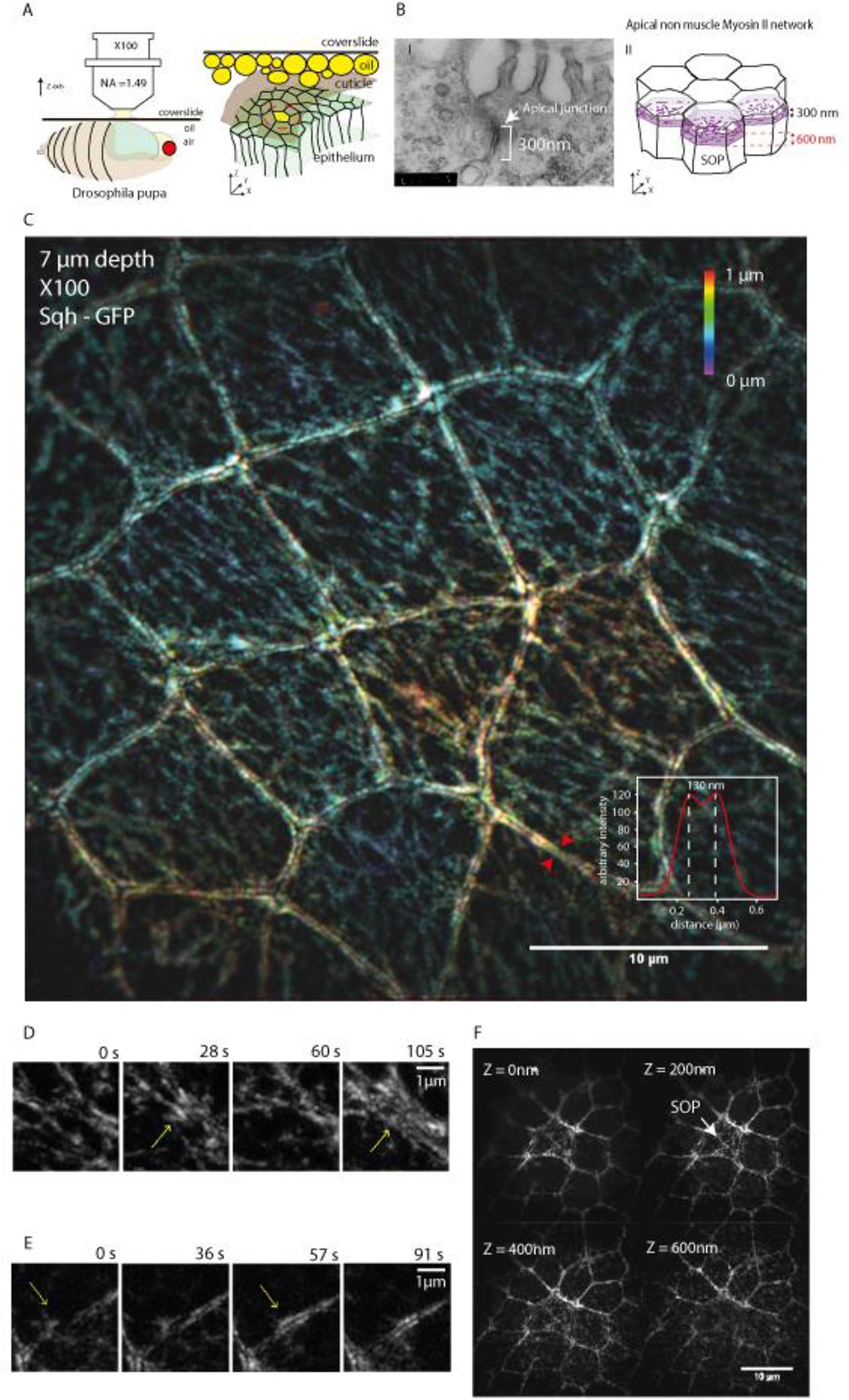
3D RIM Myosin II imaging in live *Drosophila* notum. (A) Cartoon depicting the living pupa with pupal case (brown) removed on the notum (green) and experimental conditions during acquisition and cartoon depicting the location of medial and junctional Myosin II (magenta) of epidermal cells of the Drosophila pupal notum. The yellow cell corresponds to a Sensory Organ Precursor cell (SOP). The cells inside the dotted red circle are imaged in (C). *(B) (I)* TEM picture shows an orthogonal section of the junctional domain separating two epidermal cells of the pupal notum. The apical surface exhibits the typical organisation of microvilli. The electron dense zone at the cell-cell boundary corresponding to the adherent junction (arrow) has a thickness of 300 nm (white brackets). *(II)* Cartoon depicting the mosaïc of the pupal notum cells composed of epidermal cells and of sensory organ precursors (SOP). The medial Myosin II network of SOP is positioned slightly basally relative to that of epidermal cells. (C) RIM 3D widefield view (made of four slices 200 nm apart) of the medial and junctional Myosin II network at the apical plane as described in (A) (Movie S8). The fluorescence intensity plot between the two red arrowsheads at the level of junctional Myosin II networks of two adjacent cells revealed a 130 nm resolution which is maintained constant throughout the whole field of view. (D) Time-lapse imaging of Myosin II (see also Movie S8) showing the spatial reorganization (yellow arrow) and contractile behavior of medial Myosin II. (E) Fast reshaping of junctional Myosin with recruitment of Myosin II filaments (yellow arrow). (F) Four consecutive RIM slices at the level of the medial Myosin II network denoting the high axial resolution enabling to discriminate the thickness and positioning of the medial Myosin network in SOP about 300 nm basally relative to that of its neighboring epidermal cells.

In Figure 6C, we noted that the 3D image reconstruction of medial Myosin II network was rather uniformly colored. This observation suggested that apical myosin is spatially restricted to a small section (about 300 nm thick) of the epidermal cells (see the color scale bar). In line with this conclusion, transmission electron microscopy images on sections along the apical-basal axis, showed that the thickness of the adhesive cell-cell contact, was of only 300 nm (Figure 6B part I), which corresponds well to the Myosin II belt previously described (Ebrahim et al., 2013). RIM high axial resolution was also illustrated in Figure 6F where it elegantly allowed to discriminate the lower apical positioning and larger thickness of the Myosin II networks of sensory organ precursor cells compared to that of surrounding epidermal cells: at Z=0 nm up to Z= 600 nm Myosin medial network was well observed in SOP cells, while medial Myosin of the neighboring epidermal cells, barely detectable at Z=200 nm, was well observed at Z=400 nm and Z=600 nm. These observations, summarized in the cartoon Figure 6B, come at variance with a previous conclusion that Myosin II was mostly junctional in neighboring non-SOP cells (Couturier et al., 2017).

The above results underscored the versatility of RIM for imaging a variety of tissues from large fields of views to macromolecular motion. In the following experiment, we show that RIM can also help visualizing large scale cell movements. We chose to focus on a process of collective cell migration occurring deep in a tissue, the migration of border cells in D*rosophila* ovary labeled on F-actin by UtrABD-GFP. At stage 9, these border cells perform an invasive migration on the intervening nurse cells to finally reach the oocyte at stage 10. A 75 min movie (Movie S9) shows, with a constant image quality, the migration of border cells from the anterior epithelial surface to the center, 60 microns deep, of the egg chamber. In addition to the super-resolved reconstructions, classical widefield images were obtained by simply summing the speckle frames as proposed in (Mudry et al, 2012). The widefield images, which resemble transmission-microscopy images thanks to the out-of-focus fluorescence (Shain et al., 2017), permit to locate the migrating cells in its complex environment (Figure 7A). The migration process was clearly unaffected by the 200,000 speckles illumination. Remarkably, a constant transverse resolution was obtained, whatever the position of the cells. At the end of the migration, close to the oocyte, the resolution was still about 160 nm (Figure 7B). Altogether, these data demonstrate the ability of RIM to image deep inside living tissues, with high resolution and no apparent toxicity.

**Figure 7.**
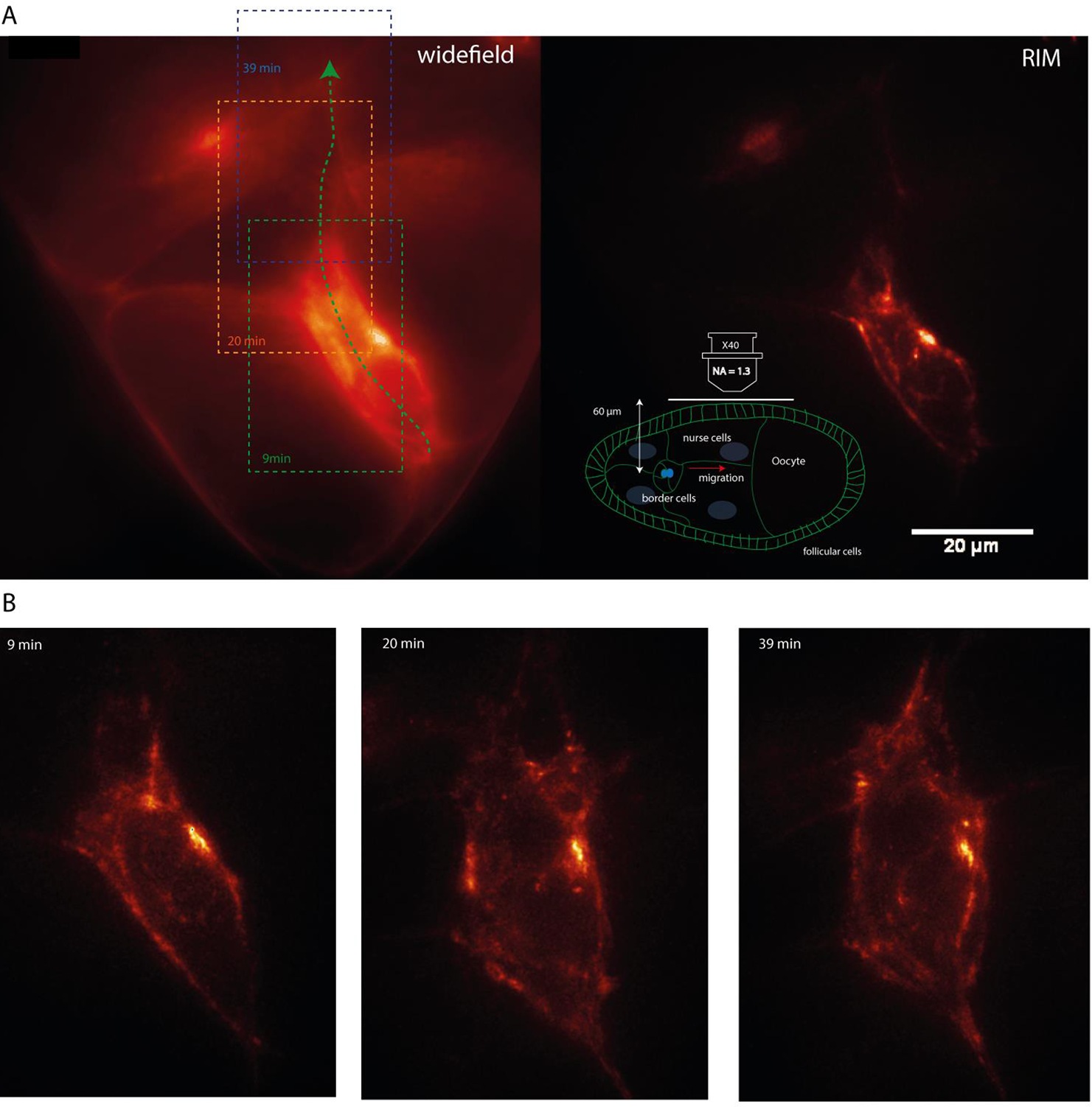
**Border cell migration in live Drosophila ovary.** (A) 3D RIM imaging of F-actin labeled by UtrABD-GFP in *Drosophila* ovary during the detachment process of migrating border cells. The inset shows the experimental conditions during acquisition and the border cell migration trajectory. The 3D images are made of 10 different focal planes with axial step of 1 µm. 200 speckles images are recorded per slice. Left (widefield): The deconvolved speckle images are summed to form a widefield image. The latter permits to locate the migrating cells in their environment as the out-of-focus fluorescence provides a transmission-microscope-like image. Right (RIM): a super-resolved reconstruction is obtained using algoRIM to gain details on the actin network. Sole the maximum intensity from the 10 slices is presented. The trajectory of the migration (the dotted green arrow and the red arrow in the ovary scheme) indicates the position of the collectively migrating cells at 9, 20 and 39 min. (B) Images of the migrating cells taken 50 μm deep inside the ovary at 9, 20 and 39 min following the trajectory shown in (A). The microfilaments of actin are well resolved during this collective cell migration.

## Discussion

RIM is a simple live-cell imaging technique, based on speckle illumination of the samples, that combines super-resolution, robustness to aberration and scattering, low toxicity, and good temporal resolution which, altogether, makes it particularly suited for imaging intracellular dynamics from molecular motion to collective cell migration in thick specimens.

### Simplicity of implementation and ease of use of RIM

The first asset of RIM is the simplicity of its implementation. RIM provides a super-resolved reconstruction of the sample from a set of low-resolution images recorded under different uncontrolled speckles. Any widefield microscope can be adapted to RIM by replacing the lamp by a laser and introducing a diffuser on the illumination path to form the speckles. The knowledge of the illumination patterns being unnecessary, RIM tuning protocol is similar to that of classical widefield microscopy. Multicolor imaging requires only multilaser excitation and appropriate filtering as in widefield microscopy (Figures 1 and S5).

### Super-resolution and fidelity to the true fluorescence

The second asset of RIM lays in its original inversion scheme, algoRIM, which yields reconstructions true to the actual fluorescence dynamic range with a resolution of 120 nm transversally and 300 nm axially, matching that of the best 3D SIM (Figures 1, 2, S1 and S2).

The super-resolution achieved by RIM can be explained on theoretical grounds. A mathematical study has demonstrated that a twice-better super-resolved sample reconstruction could be theoretically obtained from the covariance of the speckle images provided that the Fourier supports of the speckle autocorrelation and observation point spread function are similar (Idier et al., 2018; summarized in Supplemental information). In practice, each speckle image is deconvolved using a Wiener filter to reduce the width of the point spread function and the statistic noise. Then, the variance of the speckle images is formed. Last, the fluorescence is estimated iteratively so as to minimize the distance between the rigorous model of the variance accounting for the speckle autocorrelation, and the empirical variance (see Supplemental information). This critical inversion step restores the fluorescence dynamic range and improves significantly the resolution compared to the process consisting in taking the standard deviation of the deconvolved speckle images (Taylor et al., 2018; Ventalon et al, 2007) (Figure S1E). Importantly, the reconstruction scheme does not use any regularisation except for the one needed for stabilizing the solution with respect to noise. As a result, algoRIM is successful on both dense and sparse fluorescent samples and avoids the common artefacts encountered in fluctuation or sparsity-based microscopy such as the over or underestimation of strongly or weakly labelled features (Marsh et al, 2018). Hence, as compared with the SEM image, RIM showed more details than dSTORM of the densely fluorescent podosome nodes (Figure 1C), and when the same vimentin filaments were observed by RIM or by STED microscopy, the same image was obtained at the same resolution (Figure 2B), which underscored the fidelity of RIM image reconstruction.

### Robustness to aberrations and scattering

To provide super-resolved reconstructions, the inversion procedure of RIM requires data with sufficient signal to noise ratio and a model for the point spread function and the speckle autocorrelation. Contrary to SIM, RIM is not affected by the illumination deformations induced by the sample, the lens imperfections or the experimental drifts as the speckle autocorrelation is insensitive to aberration or scattering (Goodman, 2007). Moreover, the speckle dynamic range being much larger than the dynamic range of periodic grids, RIM images are more contrasted than SIM images and less affected by the background noise as seen in Figure 1D. As a result, RIM provided super-resolved reconstructions in conditions where SIM failed (Figure S4C, MovieS6). Moreover, the transverse resolution, about 120 nm, could be maintained over large fields of view (100 µm x 100 µm) and over extended periods of time (more than an hour) (Figures 6 and 7 and Movie S9). To our knowledge, such a constancy of performance in space and time when observing thick live samples has only been obtained (though at a lower transverse resolution of about 230 nm) with a lattice light sheet microscope equipped with adaptive optics on both the illumination and observation paths to minimize specimen-induced optical aberrations (Liu et al., 2018). The unique combination of super-resolution and resistance to aberrations of RIM was demonstrated in a spectacular way by the visualization of Myosin II minifilaments motion deep inside (30 µm) a developing Drosophila leg (Figure 5). Previously, minifilaments had only been observed in cultured cells, never in thick living tissues (Fenix et al., 2016; Hu et al., 2017).

### Low toxicity

RIM phototoxicity level was clearly compatible with functional imaging: it did not detectably affect the podosomes reorganization during 20 min recording time (Figure 3), nor the collective migration of border cells during 75 min (Figure 7, Movie S9). Most remarkably, RIM did not alter the highly stress-sensitive mitosis duration in *S. pombe* when imaging kinetochore motion from prophase to telophase (Figure 3).

### Temporal resolution

Temporal resolution is a weak point of all super-resolution microscopy methods, especially when large fields of view are required. In this work, the temporal resolution of RIM was essentially limited by the read-out time and electronic noise of the camera. An important feature of RIM is that the number of speckle images used for forming one super-resolved image can be adapted to the sample dynamics after the recording: from the same data set, corresponding to thousands of speckle frames recorded every 12 ms, one can derive different movies of increasing temporal resolution, though at the cost of a deteriorated illumination homogeneity. In practice, 150 speckles were sufficient to observe the sparse and fast moving PCNA (Figure 3C). In addition, the exchangeable role of each individual speckle frame makes RIM particularly well adapted to interleaved reconstruction. Hence, shifting sets of 800 speckles by 10 speckles provided a seemingly continuous sliding of myosin molecules at a temporal resolution of 120 ms (Movie S8).

### Which perspectives?

RIM combines the key advantages of SIM such as super-resolution, low toxicity, good temporal resolution, no need for specific fluorophores, but with the unrivalled ease of use of widefield fluorescence microscopy and the ability to image deep into samples without sophisticated adaptive optics. Of course, there is room for improvements. Hence, faster cameras with continuously improving data acquisition rate, multifocus techniques to record several planes of the sample simultaneously (Abrahamsson et al., 2013) as well as the use of complementary speckle sequences (Gateau et al., 2017) for reaching the illumination homogeneity faster, should ameliorate the temporal resolution. The development of fluorophores that emit at wavelengths much larger than the excitation should alleviate the deformation of the observation point spread function to probe even deeper into the sample. There is also the possibility to modify the nature of the speckle and of the excitation to further improve the resolution (Labouesse et al., 2017, Negash et al. 2018).

To conclude, we believe that RIM will fill the expectations of cell biology laboratories, in line with the growing need for simple, fast, super-resolved functional imaging of live-cells within normal or pathological tissues or model organisms. It is worth noting that the mathematical concepts of RIM applies to all imaging techniques in which the recorded data are linearly linked to the sought parameter times an excitation field. Ultrasound imaging, diffraction microscopy, microwave scanning, photo-acoustic imaging among others, could benefit from the philosophy of this novel approach.

## Acknowledgements

We thank François Payre, Paul Mangeat and Malek Djabali for proofreading the manuscript. We thank the Imaging Core TRI and Drosophila facilities of the CBI, and Isabelle Fourqueaux (CMEAB) for SEM imaging. We thank INSERM Plan Cancer 2014-2019 and Toulouse Cancer Foundation for partial financial support. We thank David Villa for video editing (https://www.scienceimage.fr/).

## Author contributions

T.M., S.L., M.A., J.I. and A.S. conceived the project. A.S., M.A., S.L., and J.I. elaborated the theoretical RIM concept. S.L. developed and implemented the reconstruction method algoRIM. T.M. adapted algoRIM to the experimental dataset and made all RIM reconstructions, figures and movies. T.M. and T.L. implemented and automatized the RIM experiment. T.M. analysed the data with the help of all other authors. R.P., and A.B. prepared macrophages and provided the SEM images of podosomes and the high-density STORM raw images which were processed by T.M. G.M. prepared the C. *elegans* and provided its EM image. E.M. and T.M. designed the *Drosophila* leg experiments using SIM, RIM and Airyscan. M.S., R.L.B., and X.W. supervised the molecular stainings of non-muscle Myosin II in the *Drosophila*. T.M., M.P., and R.L.B. did the RIM and EM imaging of the *Drosophila* pupal notum. E.V. provided the SIM images from Elyra Zeiss. S.A., and C.R. performed the STED images of vimentin in HUVEC cells. S.C., N.C. and E.C. provided the 3D PALM images of *S. pneunomia*. A.G. made the U2OS cell lineage and PCNA staining for RIM live imaging. C.R. made the *S. pombe* cell lineage for kinetochore RIM imaging. X.W. supervised the strain and sample preparation of *Drosophila* ovary for border cells migration experiments. A.S. and T.M. wrote the manuscript. All authors discussed the results and commented on the manuscript.

## Declaration of interests

AlgoRIM described herein is covered by a provisional invention statement filed by S.L., T.M., J.I., M.A., and A.S. and covered by the CNRS.

## Movie Captions

Movie S1: Comparison between the raw images in dSTORM and RIM, related to Figure 1. Movie S2: 3D RIM reconstruction of fixed Vimentin filaments, related to Figure 2.

Movie S3: Dynamics of F-actin podosomes, related to Figure 3. Movie S4: Dynamics of PCNA, related to Figure 3.

Movie S5: 3D RIM reconstruction of *S. pombe* kinetochores compared to the 3D widefield image, related to Figure 3.

Movie S6: Comparison of RIM and SIM raw images in the fixed *Drosophila* leg, related to Figure 4.

Movie S7: Comparison of the RIM and Airyscan reconstructed images in the fixed *Drosophila* leg, related to Figure 4.

Movie S8: Dynamics of the live myosin network in the *Drosophila* leg (at the apical and basal plane), notum (apical plane and SOP) and egg chamber (basal plane), related to Figure 5, S5 and 6.

Movie S9: Migration of the border cells in *Drosophila* ovary, comparison between RIM and widefield images, related to Figure 7.

---

**Figure S1:**
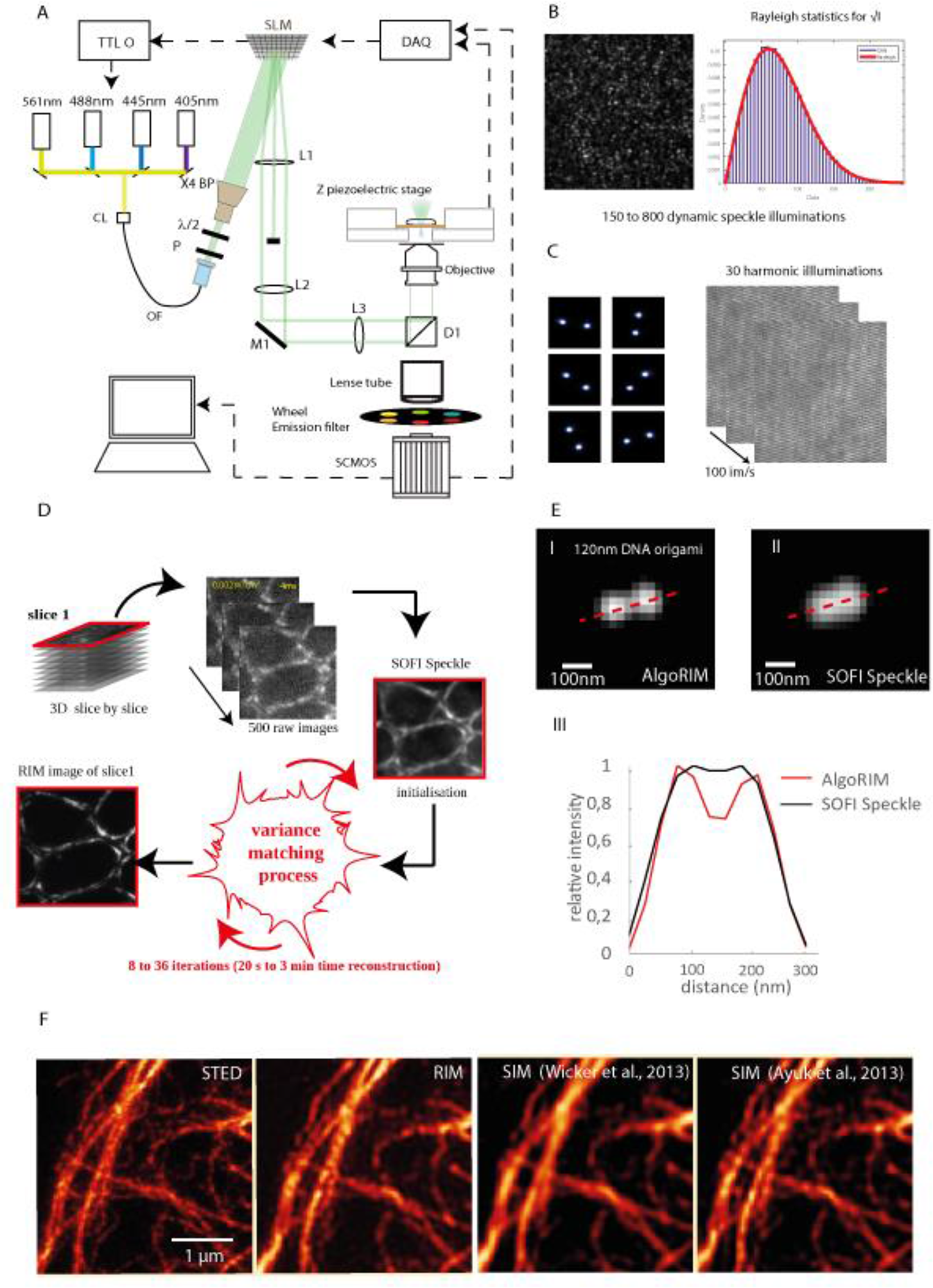
R**I**M **experimental and reconstruction principles, related to** Figure 1. (A) Detailed RIM optical setup. Four fast diode lasers are used to illuminate the sample. The light is shaped into a 8 mm collimated TEM_00_ beam thanks to an apochromatic fiber collimator (FC), an optical fiber (OF) and a beam expander (BP). After adjusting its polarization, it illuminates a fast Spatial Light phase binary Modulator (SLM), which is conjugated to the object plane of the microscope *via* the relay lenses L1-3. In the intermediate Fourier plane, a quarter wave plate, which is replaced by a pizza laminated polarizer in the SIM configuration, is introduced to produce a circular polarization state for the speckle. Aquadrichroic beam splitter with a cut off centered at 405/488/561/633 nm (D1) reflects the laser beam towards the inverted microscope. The fluorescence is collected *via* the tube lens on a SCMOS camera after appropriate filtering using a wheel filter. A Z-piezoelectric stage permits to translate the sample through the focal plane. (B) Speckle intensity obtained with a binary phase SLM. The square root of the illumination fits the Rayleigh distribution statistics. (C) The same SLM can be used to provide the periodic illuminations used in SIM. Intensity obtained at the pupil or object plane of the microscope. (D) Principle of RIM. The fluorescence density of the sample is estimated by forming the variance of the deconvolved speckle images and estimating the sample so that the variance model best matches the experimental one (see RIM theory in the Supplemental Information). The key point of RIM is that the variance model does not assume that the speckle correlation is a Dirac function. The estimation is performed using an iterative conjugate gradient scheme. This procedure ensures a linear link between the reconstruction and the sample fluorescence density and improves significantly the resolution compared to the SOFI-speckle approach, which consists in taking the standard variation of the deconvolved speckle images (Taylor et al, 2018; Ventalon et al, 2007). (E) Low-resolution images of DNA nanoruler (SIM 120 B) are obtained for 200 different speckles. (I) Super-resolved image of the sample using AlgoRIM; (II) super-resolved image using the square root of the variance; (III) intensity profile plot along the dashed line in (I) and (II). (F) From left to right, STED, RIM and SIM reconstructions of the same vimentin network from fixed HUVEC cell using a fluorescence antibody dedicated for STED microscopy with excitation at 561nm and emission at 700 nm (large Stokes shift). The RIM reconstruction is very close to the STED image. SIM reconstruction performed using the algorithm of Wicker et al., (2013), implemented in the Zeiss Elyra, in which the period and phases of the periodic pattern are recovered from an analysis of the low-resolution images, did not give satisfactory results. Better results were obtained by resorting to a more sophisticated algorithm named filtered blind-SIM (Ayuk et al., 2013), that did not assume the periodicity of the illumination but requires significantly more computational time than the classical approach. This example points out the robustness of RIM processing by comparison with SIM.

**Figure S3.**
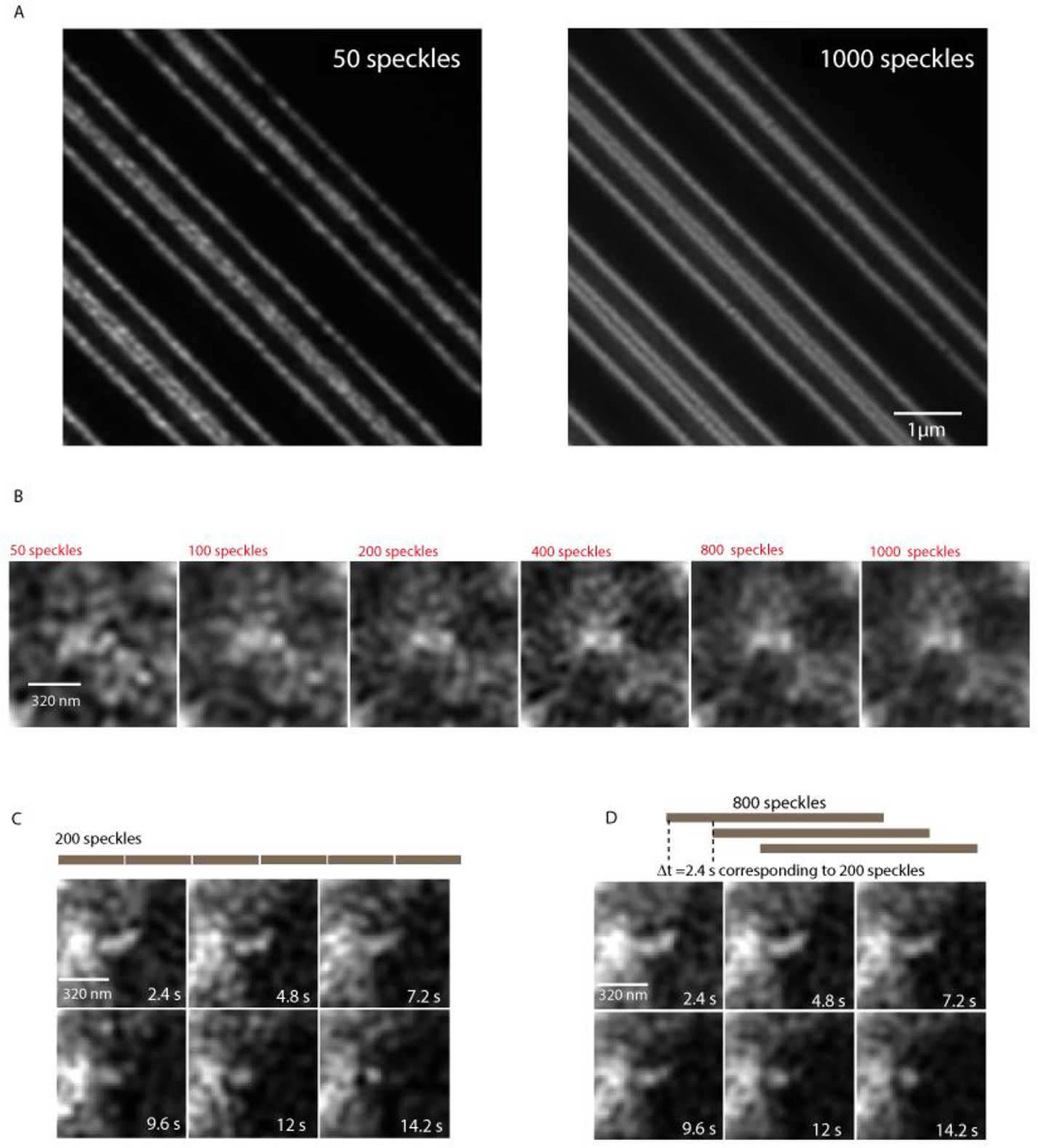
: Influence of the number of speckles in the RIM reconstruction, related to. Figure 3. (A) RIM reconstruction from 50 or 1000 speckle images of a calibrated sample (ARGO-SIM Argoligth). Interdistance between the two center lines from bottom left to top right sold for being 120-90-60 nm but estimated to be 140, 120 and 90 nm, respectively, using the calibrated pixels of the image. RIM resolution is shown to distinguish the lines of the middle pattern. 1000 speckles yield a more homogeneous image of the sample. (B) RIM reconstruction of a podosome core *versus* the number of speckle images used for the data processing. Extracted from Movie S3. Scale bar 800 nm. Due to the continuous spatial reorganization of podosomes, 400 speckles is a better choice than 1000 speckles. (C) RIM images sequences of F-actin clusters with classical RIM reconstruction using 200 speckles (12 ms per speckle). Scale bar, 800 nm. (D) RIM images sequences of F-actin podosomes core with interleaved reconstruction: 800 speckles shifted by 200 speckles are used for each time point. Scale bar, 800 nm.

**Figure S4:**
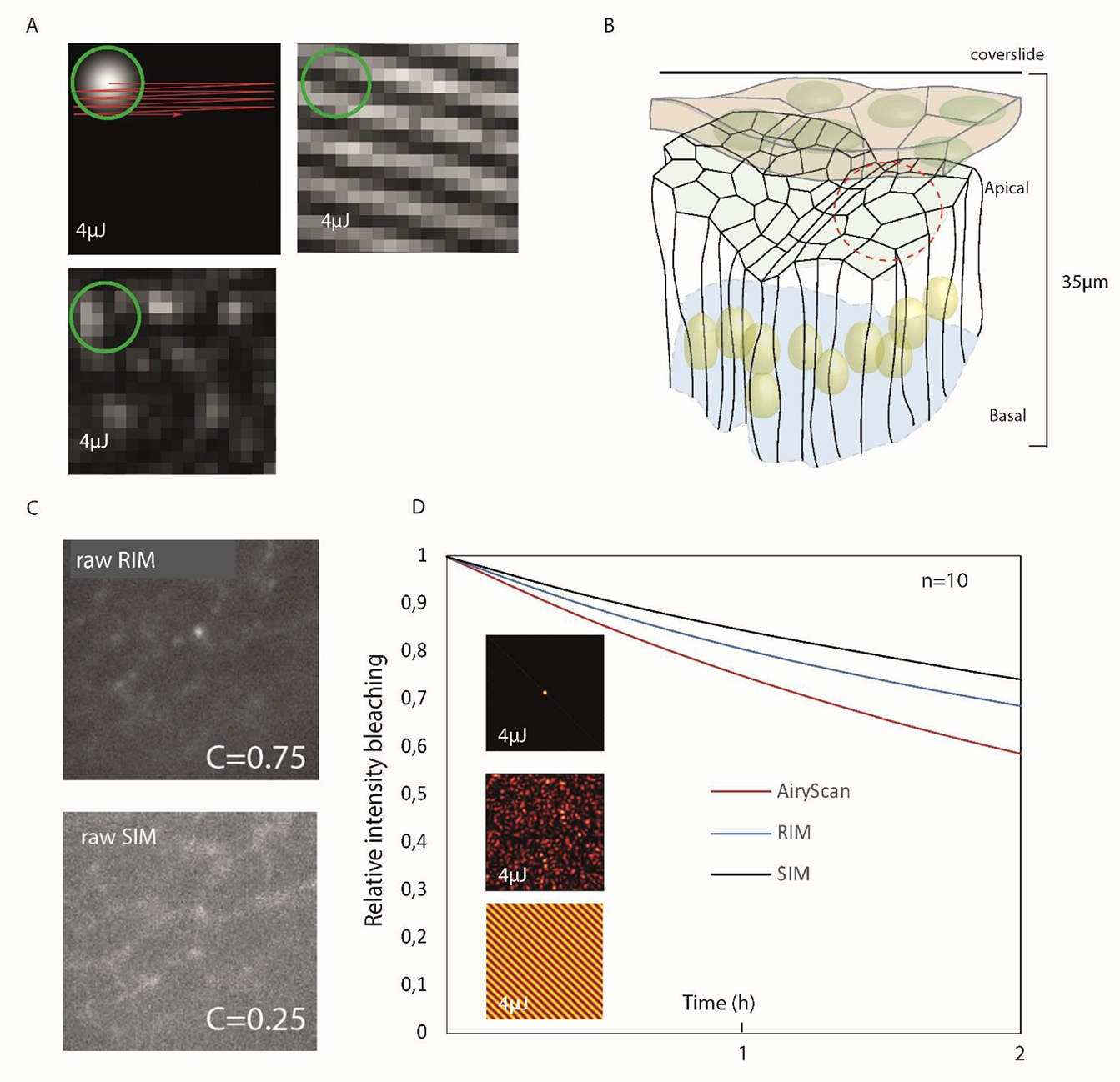
Imaging of the Myosin network in Drosophila leg, bleaching and robustness to aberrations, related to Figure 4. (A) Illustration of the imaging conditions using AiryScan, RIM and SIM. In Airyscan (top left) the beam is focused at the object plane and translated. In SIM (top right) the illumination is a periodic grid which is translated and rotated. In RIM (bottom left) the illumination correspond to hundreds of different speckles. All the experiments are conducted by injecting the same total energy of 4 µJ per diffraction limited pixel, indicated by the green circle. (B) Description of the bilayer epithelium of *Drosophila* leg. The peripodal epithelium layer in magenta introduces a first optical layer. The apical part of the second epithelium layer is the area imaged by AiryScan, SIM and RIM (indicated by the dashed red circle). (C) One raw RIM image and one raw SIM image of the leg epithelium at 7 µm depth. The periodic grid of SIM is not visible and the SIM image contrast C=(Imax-Imin)/(Imax+Imin) =0.25 is smaller than that obtained by RIM, C=0.75. In this configuration, SIM reconstruction with SIMcheck (Ball et al., 2015) fails. (D) Decay of the mean fluorescence intensity (bleaching) as a function of time averaged over n=10 different fields of view for the three imaging modalities, Airyscan, SIM and RIM.

**Figure S5.**
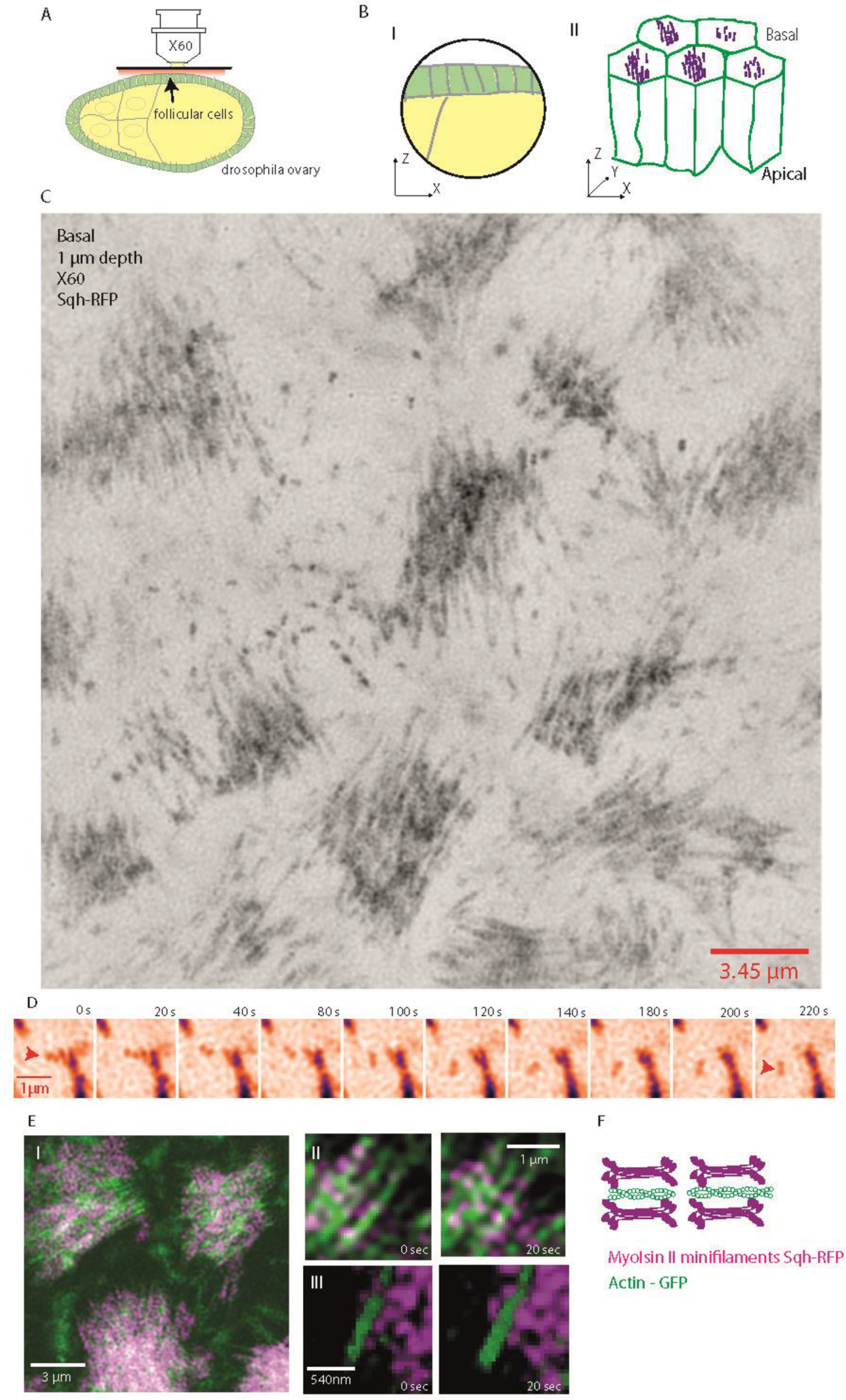
RIM imaging in live Drosophila ovary, related to. Figure 5 **and** Figure 6. (A) Experimental conditions during acquisition. The imaged plane lays about one micron deep inside the ovary. (B) *(I)* Schematical representation of follicular epithelial cells (FEC) of Drosophila egg chamber at stage 9. (II)At the basal pole of FEC, facing the extracellular matrix, Myosin II filaments are organized in parallel bundles lying along actin filaments. (C) Snapshot of a 3D RIM widefield view of the parallel bundles of Myosin II at the basal plane of live follicular epithelial cells (see Movie S8). (D) Zoom on the rotatory movement of a single Myosin II minifilament at the basal plane of follicular epithelial cells (red arrowhead). (E) (I) Two-color live-imaging of Myosin II-RFP (Sqh-RFP, magenta) together with Actin labelled with the Actin Binding Domain of Utotrophin tagged with GFP (Utrophin-ABD-GFP, green) at the basal surface of follicular epithelial cells discloses the alignment of Myosin II minifilaments with actin filaments, cartooned in (F), showing the ability to monitor the dynamics of two fluorescent probes with a high spatio-temporal resolution. *(II)* and *(III)* High magnifications.

