## Supplemental Information for "Super-resolved live-cell imaging using Random Illumination Microscopy"

**1. RIM theory and data processing**

In this supplementary part, we explicit how the stack of low-resolution speckle images is processed in order to form one super-resolved image reconstruction of the sample. We assume that the object of interest is the thin slice of sample that is located at the focal plane of the microscope. The fluorescence stemming from markers outside the focal plane is considered as noise. The fluorescence density of the sample slice is denoted by f(**r**) where **r** is a two-dimensional vector that indicates a transverse position at the focal plane. The 3D reconstruction is performed by imaging different slices of the sample via a translation through the focal plane. We first summarize the mathematical demonstration of the super-resolution capacity of RIM (Idier et al., 2018) and compare it to that of fluctuation microscopy. Then we detail the data processing that has been implemented to reach a two-fold resolution gain.

*1.1 Modeling RIM data*

The 2D intensity I of the low-resolution speckle image at the observation point $\mathbf{r}_{obs}$ on the camera is a random variable that can be modeled as,

$$I(\mathbf{r_{obs}})= \int f(\mathbf{r})\, E(\mathbf{r})\, H_{\lambda'}(\mathbf{r_{obs}}-\mathbf{r})\, d\mathbf{r}\ , \quad (1)$$

where f(**r**) is the fluorescence density, $H_{\lambda'}(\mathbf{r})$ is the observation point spread function (PSF) at the fluorescence wavelength λ' and E(**r**) is the speckle intensity.

In our epi-illumination microscope, the speckle and observation PSF are formed via the same objective of numerical aperture NA. The speckle can be modeled as a sum of plane waves with transverse wave vector **k** and random phase φ(**k**) which satisfy, $<\exp[i\varphi(\mathbf{k})]>=0$ and $<\exp[i\varphi(\mathbf{k})-i\varphi(\mathbf{k}')]>=\delta(\mathbf{k}-\mathbf{k}')$ where <> stands for the ensemble average. We note E(**r**) the speckle intensity at position **r** in the focal plane,

$$E(\mathbf{r}) = \left| \int l_{\lambda}\, (\mathbf{k})\, \exp[i\varphi(\mathbf{k})]\, \exp[i\mathbf{k.r}]\, d\mathbf{k} \right|^2 \quad (2)$$

where λ is the illumination wavelength, $l_{\lambda}(\mathbf{k})=1$ if $|\mathbf{k}|<2\pi NA/\lambda$ and 0 elsewhere, and scalar approximation has been used. The theoretical ensemble average of E is a constant <E> and its covariance is,

$$S(\mathbf{r},\mathbf{r'})=< [E(\mathbf{r})-<E>]\ [E(\mathbf{r'})-<E>] >. \quad (3)$$

Using Eq. (2), one shows that $S(\mathbf{r},\mathbf{r'})$ is similar to $H_{\lambda}(\mathbf{r}-\mathbf{r'})$, the point spread function of the microscope at the illumination wavelength defined as,

$$H_{\lambda}(\mathbf{r}) = \left| \int l_{\lambda}(\mathbf{k})\, \exp[i\, \mathbf{k.r}]\, d\mathbf{k} \right|^2. \quad (4)$$

The first and second moments of fully developed speckles are thus well known and they are insensitive to scattering distortion or aberration (Goodman, 2007).

*1.2 Super-resolution capacity of RIM*

To eliminate the unknown illuminations from Eq. (1), we form the covariance of the image intensity

(Idier et al., 2018),

$$C(\mathbf{r}_{obs1},\mathbf{r}_{obs2}) = \langle [I(\mathbf{r}_{obs1}) - \langle I(\mathbf{r}_{obs1})\rangle] [ I(\mathbf{r}_{obs2}) - \langle I(\mathbf{r}_{obs2})\rangle] \rangle$$

$$= \int f(\mathbf{r}_1) f(\mathbf{r}_2)\, S(\mathbf{r}_1-\mathbf{r}_2)\, H_{\lambda'}(\mathbf{r}_{obs1}-\mathbf{r}_1) H_{\lambda'}(\mathbf{r}_{obs2}-\mathbf{r}_2) d\mathbf{r}_1 d\mathbf{r}_2 \quad (5)$$

which depends on the known speckle covariance and observation PSF. The covariance being a non-negative definite operator, S and C admit unique non-negative definite square roots $S^{1/2}$and $C^{1/2}$ which satisfy, (Mercer, 1909)

$$S(\mathbf{r}_1-\mathbf{r}_2) = \int S^{1/2}(\mathbf{r}_1-\mathbf{r}')\, S^{1/2}(\mathbf{r}'-\mathbf{r}_2)\, d\mathbf{r}', \quad (6)$$

where the translational invariance of speckle properties has been used and,

$$C(\mathbf{r}_{obs1},\mathbf{r}_{obs2}) = \int C^{1/2}(\mathbf{r}_{obs1},\mathbf{r}')\, C^{1/2}(\mathbf{r}',\mathbf{r}_{obs2})\, d\mathbf{r}'. \quad (7)$$

Then, the image covariance can be written as,

$$C(\mathbf{r}_{obs1},\mathbf{r}_{obs2}) = \int C^{1/2}(\mathbf{r}_{obs1},\mathbf{r}')\, C^{1/2}(\mathbf{r}',\mathbf{r}_{obs2})\, d\mathbf{r}'$$

$$= \int d\mathbf{r}' \int f(\mathbf{r}_1) S^{1/2}(\mathbf{r}_1-\mathbf{r}')\, H_{\lambda'}(\mathbf{r}_{obs1}-\mathbf{r}_1)\, d\mathbf{r}_1 \int f(\mathbf{r}_2) S^{1/2}(\mathbf{r}'-\mathbf{r}_2)\, H_{\lambda'}(\mathbf{r}_{obs2}-\mathbf{r}_2)\, d\mathbf{r}_2 . \quad (8)$$

Now, if the Fourier support of S is equal or included in that of $H_{\lambda'}$, one can filter the speckle images so that the point spread function $H_{\lambda'}$ becomes equal to $S^{1/2}$. In this case, the operator K acting on the function q,

$$K(q) = \int q(\mathbf{r}_1)\, S^{1/2}(\mathbf{r}_1-\mathbf{r}')\, H_{\lambda'}(\mathbf{r}_{obs1}-\mathbf{r}_1)\, d\mathbf{r}_1 = \int q(\mathbf{r}_1)\, S^{1/2}(\mathbf{r}_1-\mathbf{r}')\, S^{1/2}(\mathbf{r}_{obs1}-\mathbf{r}_1)\, d\mathbf{r}_1$$

is definite and positive so it can be identified to the square root of the image covariance. Then, taking the diagonal terms, one obtains,

$$C^{1/2}(\mathbf{r},\mathbf{r}) = \int f(\mathbf{r}_2) K(\mathbf{r}-\mathbf{r}_2) d\mathbf{r}_2 \quad (9)$$

where $K(\mathbf{r}-\mathbf{r}_2) = S^{1/2}(\mathbf{r}-\mathbf{r}_2) S^{1/2}(\mathbf{r}-\mathbf{r}_2)$.

When the speckle is generated through the same objective as the observation point spread function, the Fourier support of $S^{1/2}$ is similar to that of $H_{\lambda'}$ so the support of K is twice larger than that of $H_{\lambda'}$. Thus, from a theoretical point of view, RIM image covariance bears enough information to reconstruct the fluorescence density with a resolution twice better than that of classical fluorescence microscopy.

#### *1.3 RIM versus Fluctuation Microscopy*

At this point, one can draw a link between RIM and Fluctuation Microscopy which is also based on the processing of the second (and sometimes higher) order statistics of ‘random’ images. In fluctuation microscopy, the sample is excited with an homogeneous intensity and one takes advantage of the natural fluctuation (or blinking) of the fluorescence with respect to time to record different realizations of a random imaging process. The images can be modeled using the same Eq. 1 as that used for RIM, E being now a random variable accounting for the emission fluctuation.

The fundamental difference between speckle illumination and fluctuation microscopy is that, in the first case, the random process exciting the fluorescence is spatially correlated, $S=H_{\lambda}$ while, in the second case, it is totally uncorrelated and S is a Dirac. This has a major incidence. Trivially, in fluctuation microscopy, the Fourier support of S is not included in that of $H_{\lambda'}$ so there is no mathematical proof that the second order statistics of the images can provide the fluorescence density over an enlarged Fourier domain. More precisely, the variance of fluctuation microscopy data simplifies to a simple convolution of the square of the fluorescence density with the square of the fluorescence density,

$$V_{fluctuation}(\mathbf{r})= C(\mathbf{r},\mathbf{r}) =\int f^2(\mathbf{r_1})H^2_{\lambda'}(\mathbf{r}-\mathbf{r_1})d\mathbf{r_1}. \quad (10)$$

This relationship permits to estimate the square of the fluorescence density on the Fourier support of $H^2_{\lambda'}$. Now, the knowledge of the Fourier transform of $f^2$, $FT(f^2)$, over a domain W, does not imply the knowledge of FT(f) over W. Hence, except if f is binary, the super-resolved reconstruction provided by fluctuation microscopy should be taken with caution as it depends on the fluorescence density in a non-linear way. In particular, the dynamics of the staining is generally lost. The weakly and strongly labeled features being often under and over estimated, respectively [Marsh, 2018].

*1.4 RIM data processing*

On the contrary, when the speckle covariance S has a bounded Fourier support, as in RIM, the fluorescence density can theoretically be estimated on an enlarged Fourier support, without loss of the staining dynamics. However, estimating the square root of the speckle image covariance matrix is impossible in practice because of the huge size of the operator. A more tractable technique consists in developing a marginal inversion procedure (Idier et al., 2018) in which the fluorescence density is estimated so as to minimize the distance between the recorded image covariance and the simulated one. Yet, while efficient, this approach cannot be applied to large fields of view as it is computationally demanding, the covariance operator scaling as MxM where M is the number of pixels on the camera.

In this work, we propose a simplified approach in which the fluorescence density is estimated so as to minimize the distance between the variance model C(**r**,**r**) and the empirical variance. Note that the covariance C(**r**,**r'**) is maximal at **r'**=**r** and tends towards 0 when **r'** moves away from **r** with a typical decorrelation length corresponding to the width of the observation Point Spread Function (PSF). Thus, most of the information on the fluorescence density is contained about the diagonal of the covariance.

Once N different speckle images $I_{l=1...N}$ of the sample have been recorded, the data processing comprises three steps :

1) Each speckle image is deconvolved with a Wiener filter. More precisely, we calculate $i_l$ so that,

$$FT(i_l) (\mathbf{k}) =FT(I_l)(\mathbf{k}) \times FT(h)(\mathbf{k}) /[|FT(H)(\mathbf{k}) |^2 + e] \quad (11)$$

where FT(g) is the two-dimensional Fourier transform of g and e is a parameter (Tikhonov regularization) which depends on the noise level of the image.

The deconvolved image $i_l$ can be written as,

$$i_l (\mathbf{r})= \int f(\mathbf{r}) E_l(\mathbf{r}) h(\mathbf{r_{obs}}-\mathbf{r})d\mathbf{r} \quad (12)$$

where we have introduced a novel point spread function, h, such as

$FT_{2D}(h)\,(\mathbf{k}) = FT_{2D}(H)(\mathbf{k})\,/[|FT_{2D}(H)(\mathbf{k})+e|^2$ .

The width of this novel PSF being smaller, the information contained by the covariance of the deconvolved speckle images is further concentrated about the diagonal, i.e. the variance.

2) Second, we estimate the empirical variance of $i_l$ as

$v(\mathbf{r}) = \sum_{l=1...N} \; [i_l\,(\mathbf{r}) - \underline{i}(\mathbf{r})\,]^2\,/N$ where

$$\underline{i}(\mathbf{r}) = \sum_{l=1...L} i_l(\mathbf{r})/N \qquad (13)$$

3) Third, we estimate f so as to minimize the cost functional,

$$G(f)= \int |v(\mathbf{r})-V(f,\mathbf{r})|^2\, d\mathbf{r} \,/ \int |v(\mathbf{r})|^2 d\mathbf{r} \qquad (14)$$

$$\text{where } V(f,\mathbf{r})= C(\mathbf{r},\mathbf{r}) =\int f(\mathbf{r_1})f(\mathbf{r_2})\; S(\mathbf{r_1}-\mathbf{r_2})\; h(\mathbf{r}-\mathbf{r_1})h(\mathbf{r}-\mathbf{r_2})d\mathbf{r_1}d\mathbf{r_2}. \qquad (15)$$

To minimize G, one needs a solver able to calculate V for a given estimation of the fluorescence f. Now, the calculation of V involves a time consuming quadruple integral (15). To speed up the iterative inversion procedure, we developed an approximate expression of V which necessitates only a double integral. To this aim, we recall that functions of two variables can be written as a series of the product of functions of one variable only. Let us introduce the bivariate real function,

$T(\mathbf{r}_1\,,\mathbf{r}_2) = S(\mathbf{r}_1-\mathbf{r}_2)\; h(-\mathbf{r}_1)h(-\mathbf{r}_2)$.

Being a non-negative definite kernel, T admits an eigenvalue decomposition as,

$$T(\mathbf{r_1},\mathbf{r_2}) =\sum_k U_k(\,\mathbf{r_1})U_k(\,\mathbf{r_2}). \qquad (17)$$

In practice, T is discretized over the pixels of the camera and implemented as an Hermitian matrix. Its singular value decomposition yields the eigenvectors of decreasing norm $U_k$. Then, the decomposition is stopped at the Kth order, (K about 10 is usually sufficient to have an accurate estimation of T). We thus obtain a limited rank approximation of T. Other limited-rank approximations are possible, but the one we choose reaches an optimal trade-off according to the Eckart–Young theorem (Eckart and Young, 1936).

The variance is then calculated using,

$$V(f,\mathbf{r})\approx\sum_{k=1...P} [w_k\,(f,\mathbf{r})]^2 \qquad (18)$$

$$\text{with } w_k\,(f,\mathbf{r})=\int f(\mathbf{r_1})U_k(\,\mathbf{r_1}\,-\mathbf{r}\,)d\mathbf{r_1}.$$

(19)

It is worth noting that decomposing $T(\mathbf{r_1},\mathbf{r_2})$ is much more efficient than decomposing $S(\mathbf{r_1}-\mathbf{r_2})$. Indeed, we have found that the number of eigenvectors required for approximating T is significantly smaller than that required for approximating S with the same accuracy.

The minimization of G is performed using a standard gradient descent algorithm (Bertero,1998). At each iteration, $f(\mathbf{r})$ is modified following,

$f^n(\mathbf{r}) = f^{n-1}(\mathbf{r}) + \alpha g^n(\mathbf{r})$

where,

$g^n(\mathbf{r}) = -2 \int [\sum_{k=1...K} w_k (f^{n-1},\mathbf{r_1}) U_k( \mathbf{r} - \mathbf{r_1} ) ] b(\mathbf{r_1}) d\mathbf{r_1}$ with $b(\mathbf{r_1}) = v(\mathbf{r_1})-V(f^{n-1},\mathbf{r_1})$ is the residue and $\alpha$ is the scalar that minimizes the fourth order polynom $g(\alpha)= G[ f^{n-1} + \alpha g^n ]$ (calculated analytically). We use an early-stopping of the iterative process as a simple (but efficient) regularization technique that basically acts as a Tikhonov regularization (Bertero, 1998).

### 2. Experimental RIM/SIM setup and acquisition software

The RIM setup is illustrated in Figure S1A. A Fiber Laser Combiner with 4 fast diode lasers (Oxxius) with respective wavelength 405 nm(LBX-405-180-CSB,) 454 nm (LBX-445-100-CSB), 488 nm (LBX-488-200-CSB), and 561 nm (LMX-561L-200-COL) is used to illuminate the sample. The diode lasers have nanosecond time response for high speed triggering and an Acousto-Optic Modulator is used to modulate the intensity of the solid state 561 nm laser. An apochromatically corrected fiber collimator (RGBV Fiber Collimators 60FC Sukhamburg) produces a collimated $TEM_{00}$ 2.2 mm diameter output beam for all wavelengths. The incident beam is rotated with an angle of 5 degrees before hitting a X4 Beam Expander beam (GBE04-A) to be shaped into a 8.8 mm $TEM_{00}$ beam which illuminates, with the appropriate polarization, a fast Spatial Light phase binary Modulator (SLM : QXGA Fourth Dimensions). The latter is conjugated to the image plane and generates different speckle patterns by displaying masks with randomly independent 0 or $\pi$ phases at each pixel. The relay lens L1 is an achromatic doublet with visible reflective coating (Thorlabs AC254-500-A). In the intermediate Fourier plane of the illumination path we introduce a quarter wave plate to produce a circular polarization state for the speckle. We also block the zero order of the SLM. A second relay lens system L2 (Thorlabs AC254-150-A) and L3 (Thorlabs AC254-135-A) is used to adapt the beam to all objective lenses. A Quadrichroic Beamsplitter with a cut off centered at 405/488/561/633 nm (Di01-R405/488/561/635-25x36 Semrock) reflects the laser beam towards the inverted microscope (Tei Nikon).

The same setup has been used for providing 2D SIM images. In this case, the SLM displays a periodic pattern (Figure S1C). At the intermediate Fourier plane of the illumination path, a Fourier mask is introduced to reject the undesired spots from the SLM pixelisation and a pizza laminated polarizer with 12 segments of tangential linear polarization is used to obtain the required linear polarization of the two beams for each azimutal angle (colorPol® VIS 500 BC3 CODIX). An insensitive polarization Dual Band Dichroic Mirror is also used to transmit both 561nm and 488nm harmonic patterns (Di01-R488/561-25x36 Semrock).

Three different objective lenses have been used in the experiments and are detailed in Table 1. Note that, contrary to the SIM mode, changing the objectives or the wavelengths for the RIM mode did not require any change in the SLM masks or additional tunings.

The fluorescence is collected via the objective and tube lens onto a SCMOS camera after appropriate filtering. Three band pass filters are used : for the green fluorescence protein (GFP) emission : Semrock FF01-514/30-25, for mcherry protein : FF01-609/54-25 SEMROCK, and finally for the fluorophore used in correlative STED imaging a band pass filter centered around 670 nm (FF01-676/29-25 SEMROCK). A motorized high-speed wheel filter is used to sequentially turn the two band pass filters within 30 ms after the 200 speckle frames or the 30 SIM frames.

For forming the 3D image, the sample is translated through the focal plane with a piezoelectric Z stage (Z INZERT PIEZOCONCEPT).

To synchronize the illumination and data recording, the Rolling Shutter output of the SCMOS camera is used to trigger the mask change on the SLM. Then, the SLM output triggers the laser when the binary phase mask is stable. For SIM, the protocol and Arduino program are described in (Förster et al, 2015). A script from micromanager software was written to select the number of speckles used, the acquisition time for each speckle-image, the number of planes along the Z-axis, their interdistance and the number of colors used for the acquisition.

### 3. RIM acquisition parameters

**Table 1** : Wavelengths of excitation and emission and objective characteristics for RIM experiments.

| Sample | $\lambda_{ex}$ (nm) | $\lambda_{em}$ (nm) | NA | Lenses |
|---|---|---|---|---|
| GATTA 120B | 488 | 514 | 1.49 | CFI SR APO 100XH ON 1,49 DT 0,12 NIKON |
| GATTA 140YBY | 488 | 514 | 1.49 | CFI SR APO 100XH ON 1,49 DT 0,12 NIKON |
| | 561 | 609 | 1.49 | CFI SR APO 100XH ON 1,49 DT 0,12 NIKON |
| HUVEC labeled for vimentin | 561 | 700 | 1.49 | CFI PLAN APO LBDA 60XH 1,4/0,13 NIKON (Figure 2)<br><br>CFI SR APO 100XH ON 1,49 DT 0,12 NIKON ( Figure S1) |
| *S. pneumoniae* | 488 | 514 | 1.49 | CFI SR APO 100XH ON 1,49 DT 0,12 NIKON |
| Live and fixed podosomes | 488 | 514 | 1.49 | CFI SR APO 100XH ON 1,49 DT 0,12 NIKON |
| PCNA UO2S cells | 488 | 514 | 1.49 | CFI SR APO 100XH ON 1,49 DT 0,12 NIKON |
| *S. pombe* kinetochore | 561 | 609 | 1.49 | CFI SR APO 100XH ON 1,49 DT 0,12 NIKON |
| *C.elegans* intestine | 488 | 514 | 1.49 | CFI SR APO 100XH ON 1,49 DT 0,12 NIKON |
| Fixed and live *Drosophila* leg | 561 | 609 | 1.4 | CFI PLAN APO LBDA 60XH 1,4/0,13 NIKON |
| live *Drosophila* egg chamber | 561<br>488 | 609<br>514 | 1.4 | CFI PLAN APO LBDA 60XH 1,4/0,13 NIKON |
| Live *Drosophila* pupa | 488 | 514 | 1.49 | CFI SR APO 100XH ON 1,49 DT 0,12 NIKON |
| Live *Drosophila* ovary | 488 | 514 | 1.3 | CFI Super Fluor 40X Oil NIKON |

**Table 2** : Illumination properties, camera integration time (stream speed), axial excursion, number of speckles for RIM experiments

| Sample | stream speed | speckles/slice | $kW.cm^{-2}$/ slice | z step | z range | Total recording time |
|---|---|---|---|---|---|---|
| GATTA SIM 120B | 24 ms | 200 | 1.6 | 0 | 0 | 4.8 s |
| GATTA SIM 140 YBY | 48 ms | 200 | 2.6 | 0 | 0 | 8.8 s (red ) |
| | 24 ms | 200 | 1.6 | 0 | 0 | 4.8 s (green) |
| Live podosomes | 12 ms | 800 | 0.8 | 150 nm | 1.2 µm | 20 min |
| Fixed podosomes | 2 ms | 800 | 0.8 | 0 | 0 | 1.6 s |
| Vimentin (HUVEC) | 12 ms | 200 | 0.8 | 100 nm | 1.2 µm | 2.4 s |
| *S. pneumoniae* | 5 ms | 400 | 0.66 | 64 nm | 1.2 µm | 36 s |
| PCNA dynamics (UO2S cells) | 12 ms | 150 | 1.8 | 0 | 0 | 2 min |
| *S.pombe kinetochores* | 12 ms | 100 | 0.4 | 150 nm | 1.2 µm | 8 min |
| *C.elegans intestine* | 12 ms | 600 | 1 | 0 | 0 | 7.2 s |
| Fixed *Drosophila* leg (apical) | 12 ms | 200 | 0.8 | 150 nm | 8 µm | 28.8 s |
| Live *Drosophila* leg (apical) | 12 ms | 200 | 0.8 | 150 nm | 1050 nm | 70 s |
| Live *Drosophila* leg (basal) | 12 ms | 200 | 0.8 | 0 | 0 | 42 s |
| Live *Drosophila* pupal dorsal thorax 14 h | 12 ms | 200 | 1.8 | 200 nm | 800 nm | 329 s |
| Live *Drosophila* pupal dorsal thorax SOP 15 h | 12 ms | 200 | 1.8 | 200 nm | 800 nm | 131 s |
| Live *Drosophila* pupal dorsal thorax 15 h | 12 ms | 400 | 3.6 | 0 | 0 | 117 s |
| *Live Drosophila* egg chamber | 12 ms | 400 | 3.6 | 0 | 0 | 20 s |
| Live *Drosophila* ovary | 12 ms | 200 | 2.6 | 1 µm | 10 µm | 75 min |

### 4. RIM reconstruction method and tuning parameters

All the raw data were interpolated using zero padding with a factor 2. The excitation and collection Point Spread Functions (PSF), $H_{\lambda}$ and $H_{\lambda\square}$ were generated with the plugin PSF generator in Matlab using the Gibson Lany Model over a domain corresponding to that of the final reconstruction with a pixel size equal to 32.25 nm. The number K of eigenvectors used for estimating the variance was always taken equal to 10. The Wiener filter parameter e, the number of iterations, reconstruction time and the optional interleaved reconstruction is described in Table 3 for each sample. The numerical processing used Intel(R) Xeon(R) CPU E5-2687W v4 at 3.00GHz with 192 RAM.

**Table 3** : Parameters of the reconstruction algorithm for RIM experiments

| Sample | Wiener Filter | number of iteration | reconstruction time/image | optional interleaved reconstruction |
|---|---|---|---|---|
| Fixed podosomes | 0.0008 | 50 | 140 s | no |
| GATTA SIM 120B | 0.0018 | 14 | 36 s | no |
| GATTA SIM 140YBY | 0.005 | 16 | 44 s | no |
| | 0.0010 | 14 | 36 s | no |
| Vimentin (HUVEC) | 0.003 | 30 | 15 s | no |
| *S. pneumoniae* | 0.01 | 20 | 12 s | no |
| Live podosomes | 0.008 | 34 | 132 s | yes |
| PCNA (UO2S cells) | 0.08 | 22 | 51 s | yes |
| *S. pombe* kinetochores | 0.1 | 7 | 5 s | no |
| *C.elegans* intestine | 0.01 | 50 | 142 s | no |
| Live *Drosophila* egg chamber | 0.001 | 35 | 125 s | no |
| Live and fixed *Drosophila* leg | 0.001 | 35 | 125 s | no |
| Live *Drosophila pupa* | 0.01 | 35 | 133 s | yes |
| Live *Drosophila* ovary | 0.02 | 24 | 119 s | no |

### 5. Cell culture

*5.1 Macrophages, differentiation from primary monocytes and culture*

Human monocytes were isolated from blood of healthy donors as described previously (Van Goethem et al., 2010). Cells were re-suspended in cold PBS supplemented with 2 mM EDTA, 0.5% heat-inactivated fetal calf serum (FCS) at pH 7.4 and magnetically sorted with magnetic microbeads coated with antibodies directed against CD14 (Miltenyi Biotec). Monocytes were then seeded on glass coverslips at $1.5x10^6$ cells/well in six-well plates in RPMI 1640 (Invitrogen) without FCS. After 2 h at 37°C in humidified 5% $CO_2$ atmosphere, the medium was replaced by RPMI containing 10% FCS and 20 ng/mL of Macrophage Colony-Stimulating Factor (M-CSF) (Peprotech). For experiments, cells were used after seven days of differentiation.

*5.2 Strain construction and culture of S. pneumoniae*

Strain R4077, harboring a *ftsZ-mEos3.2* fusion at the *ftsZ* locus, was generated as follows (Bergé, 2017). Primers were designed to amplify PCR products containing: (I) the coding region of the *ftsZ* gene (1043 bp; oligonucleotides OEC11 and OEC12, and R1501 DNA as template); (II) the *mEos3.2* gene (699 bp; oligonucleotides OEC13 and OEC14), and plasmid DNA pUC57-mEos3.2 as template. PUC57-mEos3.2 contains the *mEos3.2* gene codon optimized for *S. pneumoniae* strain R6 by Genscript; and (III) the downstream region of the *ftsZ* gene (922 bp; oligonucleotides OEC15 and OEC16, and R1501 DNA as template). The PCR products were gel-purified and used as templates in a SOEing PCR using the outer primers OEC011 and OEC016. The resulting SOEing PCR product was subsequently used to transform strain R1501 . Sequences of oligonucleotides used are the following : OEC11 (caggaggtcaacctgaggttggtcgt); OEC12 (agctgatttagaaccaccaccaccagaacgatttttgaaaaatggaggtgtatcc); OEC13 (ggatacacctccatttttcaaaaatcgttctggtggtggtggttctaaatcagct); OEC14 (gttctgtattttcttttacattcatttacttaacgacgagcattatcaggcaaacca); OEC15 (tggtttgcctgataatgctcgtcgttaagtaaatgaatgtaaaagaaaatacagaac); OEC16 (cttatccgttgcacgctgactctga).

*S. pneumoniae* strains containing FtsZ-mEos3.2 (R4077) and FtsZ-GFP (R3702, fusions were used for PALM and RIM analyses, respectively. Cells were grown in C+Y medium as previously described at 37°C to an $OD_{550}$ of 0.15. Samples were collected, pelleted (3 min, 3,000 *g*) and resuspended in cold C medium. The C medium contained per liter: 5 g casein hydrolysate, 6 mg tryptophane, 11.25 mg cysteine, 2 g sodium acetate and 8.5 g $K_2HPO_4$. Cells were spotted on a microscope slide containing a slab of 1.2% agarose in C medium as described and covered with a pre-treated coverslip before imaging.

*5.3 Cell culture and maintenance for PCNA dynamics*

U20S cells stably expressing AsiSI-ER-AID, 53BP1-GFP and mcherry-PCNA (AID-DivA 53BP1-GFP mcherry-PCNA) were cultured in Dulbecco's modified Eagle's medium (DMEM Glutamax + glucose 4.5 g/L, 1 mM sodium pyruvate, Invitrogen), antibiotics (pen/strep; Invitrogen), 10% FCS (Invitrogen), 800 µg/mL geneticin (Sigma), 1 µg/mL puromycin (Invivogen), 250 µg/mL hygromycin (Invivogen) at 37°C in a humidified atmosphere with 5% CO2.

*5.4 S. pombe cell culture and strain*

Media, growth, maintenance of strains, and genetic methods were performed as previously reported (Moreno et al., 1991). Cells were grown at 25°C on yeast extract agar. Strain genotype : ST102 ndc80-GFP:: kanR cdc11-CPF::kanR ura4-D18 leu1-32 h.

*5.5 C. elegans strain and maintenance*

The *C. elegans* strain used in this study was maintained at 20°C under standard conditions as described in (Brenner, 1974). The ERM-1-GFP strain was FL378:erm-1(bab59[erm-1::mNG^3xFlag]) I.

*5.6 Drosophila strain for border cells migration*

Drosophila harboring F-actin labeled by UtrABD-GFP provided by Thomas Lecuit was used to analyze the dynamics of border cells migration.

### 6. Sample Preparation for Microscopy

*6.1 DNA nanorulers*

DNA nanoruler SIM 140 YBY has been used (GATTAquant). The mark to mark distance is 140nm between each cluster of fluorescence (Alexa Fluor® 488).

*6.2 Podosome labeling*

Macrophages plated on glass coverslips (0.17 mm ±0.005mm, Marienfeld 0117640) were unroofed and fixed as previously described (Bouissou et al., 2017) and labelled with Alexa Fluor 488-phalloidin (Molecular Probes A12379, 1/500) and Alexa Fluor 647-coupled phalloidin (Molecular Probes, A22287, 1/100) for RIM and dSTORM, respectively. Vinculin was stained using mouse anti-vinculin (clone hvin-1, Sigma-Aldrich V9131, 1/500) and a secondary F(ab')$_2$ coupled to Alexa 555 (Cell Signaling technology #4409).

For live imaging, macrophages were transduced with mCherry-LifeAct lentiviral vector 2 or 3 days before the experiments. Transduction was performed by incubating cells with lentiviral vector (MOI: 1:1) in 800 µL RMPI 1640 supplemented with 50 µg/mL protamine sulphate. After 2 h at 37°C, 1.5 mL fresh RPMI containing 10% FCS and 50 ng/mL M-CSF was added and renewed the day after transduction.

*6.3 Vimentin Staining in HUVEC cells*

For immunofluorescence microscopy, HUVEC cells were seeded and grown onto 0.17 mm H coverslip. Cells were fixed with 3.7 % PFA (Sigma Aldrich) in PBS for 10 min at 4°C, and permeabilized with 0.1 % Triton X100  for 10 min at 4°C. After washing thrice with PBS, the cells were incubated for 1 h with anti-vimentin rabbit antibody (Sigma Aldrich) and then after washing thrice with PBS, incubated with anti-rabbit Aberrior Star 635P conjugated secondary antibodies (NanoTag Biotechnologies) diluted to 1/10, and subsequently washed three times with PBS for 10 min each. Coverslips were mounted on microscope slides using home-made Moewiol-DABCO.

*6.4 PCNA labeling in U2OS cells*

AID-DivA 53BP1-GFP mcherry-PCNA stable U2OS cell line: The pmcherry-PCNA plasmid was transfected into AID-DivA 53BP1-GFP cells (Caron et al. 2015) by using the Cell Line Nucleofactor kit V (Amaxa) and selection was performed with 250 µg/mL hygromycin (Invivogen). Monoclonal, stable cell lines were selected based on PCNA-mcherry expression level.

*6.5 S. pombe kinetochore dynamics imaging*

To analyze chromosomes dynamics, kinetochore protein Ndc80 was labeled with a GFP construct (Ndc80-GFP) in wild-type cells. For live-cell analysis, the cells were put on an imaging chamber (CoverWell PCI-2.5; Grace Bio-Labs, Inc.) filled with 1 mL of 1% agarose in minimal medium and sealed with a 22 × 22 mm glass coverslip.

*6.6 C.elegans sample preparation*

L4 stages larvae were mounted on a 10% agarose pad in a suspension of 100 nm polystyrene microbeads (Polysciences Inc.) to block worm movements. For transmission electron microscopy (TEM) control L4 larvae were treated as described (Bidaud-Meynard et al, Development, 2019).

*6.7 Drosophila live and fixed leg samples*

For live imaging, imaginal leg discs from Sqh-TagRFPt$^{KI}$[3B] *Drosophila* strain were freshly dissected from prepupae (2 h after puparium formation) in Schneider medium supplemented with 15% fetal calf serum, 0.5% penicillin-streptomycin and 2 µg/mL 20-hydroxyecdysone (Sigma-Aldrich, H5142) and mounted on slides. For comparison between Airyscan and RIM, imaginal discs were fixed for 20 min in paraformaldehyde (PFA 4%) diluted in PBS. The samples were washed in PBS, re-suspended in Schneider medium and mounted on slides.

*6.8 Drosophila pupa thorax sample*

Myosin II-GFP (Daniel et al, 2018) homozygous or hemizygous *Drosophila* pupae were used in Figure 7. *Drosophila* melanogaster lines were maintained at 25°C. Pupae 15 h-17.5 h after puparium formation were prepared for live-imaging as described in (Gho et al., 1999). It consisted in removing the pupal case over the head and the dorsal thorax and in sticking the pupae on a glass side with a double-sided tape. Then, pillars made of 4 to 5 glass coverslips were positioned at the anterior and posterior side of the pupae. A glass coverslip covered with a thin film of Voltalef 10S oil was placed on top of the pillars in order to form a contact between the dorsal thorax and the glass coverslip.

*6.9 Drosophila Ovary sample*

Drosophila egg chambers were dissected and mounted in live imaging medium (Schneider's insect medium with 20% FBS) as previously described  (Prasad et al., 2007)

### 7. Other microscopy methods

*7.1 dSTORM acquisition and reconstruction (podosomes in Figure 1)*

dSTORM imaging of F-actin was performed on cells stained *in situ* by diluting Alexa Fluor 647-coupled phalloidin (Molecular Probes, A22287, 1/100) in the dSTORM buffer (Smart-kit buffer, Abbelight, France). Images were acquired using a 100X/1.49 N.A oil immersion objective (Olympus) mounted on an inverted IX83 microscope (Olympus) equipped with a TIRF module (Abbelight, France) and an sCMOS ORCA FLASH4.0 v3 (100 fps, cable camera link, Hamamatsu) camera. Samples were excited with a 640 nm (400 mW, ERROL Laser) laser at oblique illumination controlled via NEO Software (Abbelight, France). Images were collected once the density of fluorescent dye was sufficient (typically, under 1 molecule/$\mu m^2$) using an integration time of 50 ms.

The reconstruction was performed by Nanoj software SRRF. Ring radius is defined at 1.10. The interpolation is set to 3. The PSF width is estimated to be equal to 2.6 pixels. The second order statistics has been used. Drifts have been estimated with 2D cross-correlation analysis. The optional intensity weighting has been selected. 10,000 images were used for the image reconstruction.

*7.2 SIM image acquisition and reconstruction (nanorulers of Figure 1)*

The SIM acquisitions were made on the ELYRA PS.1 Carl Zeiss, with a Plan-Apochromat 63x / 1.4 Oil DIC M27 lens and a pco.edge sCMOS camera. The SIM settings using ZEN software corresponded to a periodic grid of period 23 µm for 488 nm and 28 µm for 555 nm, 3 rotations and 5 translations. The reconstruction parameters in ZEN were: SR Frequency Weighting 1, Noise Filter -6., Sectioning 100 / 83 / 83, isotropic reconstruction.

We performed the alignment of the XYZ stage, acquired the PSF for the different emission wavelengths and formed the realignment matrix between the different channels. The reconstruction and the application of the realignment matrix were realized with the acquisition software of ELYRA PS.1 (Zen Black). A global image control on the raw data and on the reconstruction was systematically done with SimCheck (Ball, 2015) .

*7.3 STED microscopy and reconstruction (vimentin filaments of Figure 2 and Figure S1)*

STED images were acquired with a Leica SP8 STED 3X microscope (Leica Microsystems, Germany) using a 100X NA:1.4 oil immersion objective. The power of the 775 nm STED laser was set to the maximum value that avoided bleaching. Then, the images were acquired with a pulsed 635 nm laser line. The parameters of the image acquisition were : 13 nm pixel size, 6 time average per line, 400 Hz scan speed. STED images were deconvolved with Huygens Professional (SVI, USA) using the CMLE algorithm, with a signal to noise ratio of 7 and 30 iterations.

*7.4 3D-PALM acquisitions (Ftz ring in Figure 2)*

PALM acquisitions were performed with an inverted microscope NIKON I-SPT microscope, equipped with the “perfect focus system” (PFS, Nikon), a piezo stage (Nano Z100-N – Mad City Labs), a TIRF objective (TIRF 100X oil ON 1.49 DIC), a quad-band dichroic mirror (97335 Nikon N-STORM TIRF Filter Set), an emission filter (ET600/50M, Chroma), an EM-CCD camera (iXon Ultra DU897, Andor), and a microscope incubator to control the temperature. The system was fully automatized and controlled with NIS-elements AR4.2 software. Excitation was controlled with an AOTF and two lasers were used for PALM acquisitions: 405 nm (Coherent Cube, 100 mW) for mEos3.2 activation and 561 nm (Coherent Sapphire, 480 mW) for mEos3.2 acquisition. The two lasers were used continuously during data acquisition (around 10,000 images with an exposure time of 30ms each). The microscope was coupled to an adaptive optics device (MicAO 3DSR – Imagine Optic). This system, thanks to a deformable mirror, allowed to minimize the optical aberrations (astigmatism, spherical, coma, trefoil, quadrafoil) and then, to introduce astigmatism for 3D single molecule localization. Aberrations corrections and astigmatism calibration were determined using fluorescent beads (FluoSpheres™ Carboxylate-Modified Microspheres, 0.2 µm, orange fluorescent (540/560), 2% solids -Invitrogen™). Z calibration, 3D particles detection and image reconstructions were performed with ThunderSTORM an ImageJ plug-in (Ovesny et al. 2014).

*7.5 Widefield microscopy (dynamics of S. pombe kinetochores in Figure 3 and Movie S5)*

Time-lapse images were taken at 25°C. Exposure times were taken at 200 ms using LED light source (SPECTRA X light engine®) reduced to 100 µW to avoid phototoxicity and photobleaching. Images were visualized with camera sCMOS flash 4 LT fitted to a microscope ((Tei Nikonwith adescribed in the Suplemental information. 100×1.49 NA objective and filters (Semrock FF01-514/30-25) for GFP. Images were recorded using the Micromanager software package (2018/03/22). Intensity and γ adjustments (threshold) were made using Image J software for better visualization with screen.

*7.6 Airyscan Imaging of Drosophila leg (Figure 4 )*

Airyscan images were acquired using a LSM880 confocal microscope with the Airyscan detector (Carl Zeiss) and equipped with a Plan-Apochromat 63x/NA 1.4 Oil M27 objective. The super resolution mode of the Airyscan was used with a calibration of 0.049 µm/pixel and a *z*-step of 0.220 µm. The solid state laser at 561 nm was used to image TagRFPt protein fused to myosin. The power density for one voxel was equal to 4 µJ (Figure S4A). Note that an automatic alignment to calibrate the Airyscan detector was done prior to the acquisition step. The reconstruction was done using the Airyscan data processing included in the ZEN software with the automatic strength (6 by default), similar to the Wiener filter used for RIM.

### 8. Data analysis

*8.1 Colocalization analysis for RIM and SIM images of GATTA SIM 140YBY nanorulers in Figure 1*

We used the Distance2maxProfile code which can be be found in the CBI ImageProcessing website at, https://imaprocess.pythonanywhere.com/Analymage/projets/.

This code plots the intensity of two channels of the image along a user's segment drawn above the clusters to be analysed and yields the interdistance between the maxima.

*8.2 Analysis of the resolution*

The Fourier ring correlation method (FRC) (Banterle et al, 2013) was used to estimate the Fourier Image REsolution (FIRE) following (Nieuwenhuizen et al, 2013).

*8.3 Diameter of confinement diffusion of PCNA*

PCNA center of mass cluster was estimated using the wavelet method implemented in ICY software (De Chaumont et al., 2012). A probabilistic method is used to track individual trajectories of PCNA cluster (Chenouard et al., 2013) The diameter clustering is the square root of the Mean Square Displacement (MSD) of each individual trajectories.
